# In Silico designed cell-penetrating anti-cancer peptide specifically inhibits VEGF-A expression

**DOI:** 10.1101/2024.02.08.579410

**Authors:** Nilanjan Banerjee, Laboni Roy, Suman Panda, Tanaya Roychowdhury, Subhrangsu Chatterjee

## Abstract

Vascular Endothelial Growth Factor-A (VEGF-A), a pluripotent cytokine and angiogenic growth factor mediates the switch to an angiogenic phenotype in cancer cells. The interaction of VEGF-A protein with the VEGF receptors (VEGFR-1and VEGFR-2) starts downstream effect that promotes angiogenesis by mediating migration and increasing the permeability of endothelial cells. A cis-regulatory elements consisting of a polypurine/polypyrimidine (pPu/pPy) tract in the proximal 36-bp region (–85 to −50), can participate in the formation of a stable higher order G-quadruplex structure (G4) which is essential for VEGF promoter activity. During cancer progression the VEGF-A G4 succumbs to cellular pressure and fails to maintain the stable structure. This shifts the balance to form duplex structure thereby increasing the rate of transcription. Earlier research has tried to develop small-molecule ligands to target and stabilize G4, however they either lack specificity or non-toxicity. Peptide on the other hand are very less studied. Here we used bioinformatics in-silico tool to develop peptides which can successfully bind and stabilize the VEGF-A G4 while reducing its gene expression. This further alters the expression fate of the VEGF-A signalling cascade and prevents angiogenesis in cancer cells. We used high resolution Nuclear magnetic resonance and molecular dynamics simulation to map the chemistry of the interaction while the qPCR and western blot allowed us to check the expression pattern of the molecules of VEGF-A signalling cascade. In this investigation, we navigate the complex interplay between peptides and quadruplex structures, unravelling valuable insights that can enhance the crafting of pharmacophores directed at the dynamic quadruplex structure. The outcomes of our study are promising, paving the way for progress in the realms of research, characterization, and optimization of peptides binding to G-quadruplexes, with potential implications for therapeutic applications.

## 1. Introduction

Vascular Endothelial Growth Factor-A (VEGF-A) plays a vital role in cancer cell development^1^. It causes increased proliferation, angiogenesis, and metastasis in a variety of tumor type^2,3^. VEGF-A, a pluripotent cytokine and angiogenic growth factor mediates the switch to an angiogenic phenotype in cancer cells. The interaction of VEGF-A protein with the VEGF receptors (VEGFR-1and VEGFR-2) starts a downstream effect that promotes formation of blood vessels by mediating migration and increasing the permeability of endothelial cells. VEGF activation is proved to be synonymous with cancer progression and hence it led to the discovery of a wide variety of anti-VEGF-based antiangiogenic drugs like bevacizumab, aflibercept, ramucirumab, sorafenib and sunitinib targeting VEGFRs^4–8^. However, several clinical trials have decoded that monotherapy or combination therapy with anti-VEGF treatment can rarely improve patient survival and adds limited benefit to patients^5,6,8,9^. All these results indicate that either drugs might not be reaching the tumor microenvironment hence it is lacking the efficiency or obstructive the VEGF-VEGFR signalling enhances compensatory mechanisms of tumour angiogenesis by augmenting the expression levels of additional angiogenic factors that these drugs do not target, circumventing the VEGF-dependent angiogenic signals. VEGF is highly overexpressed in breast cancer compared to normal or benign breast tissues. Immunohistochemistry (IHC) confirms that almost 72-90% of breast cancer is VEGF-A positive^10,11^. Adams et al. analysed several patients’ samples and proved that metastatic cancer had higher VEGF levels than normal controls^12^. Correlation has also been established between higher VEGF-A level and high histologic grade, large size, progesterone receptor negativity, estrogen receptor negativity, human epidermal growth factor receptor-2 (HER2) over-expression, and lymph node metastasis^13–15^. VEGF expression is mainly found to be regulated at transcriptional level and is modulated by variety of factors involving pH, hypoxia, inactivated tumor suppressor genes, activated oncogenes, and growth factors^16–18^. Previous study revealed the presence of a cis-regulatory elements consisting of a polypurine/polypyrimidine (pPu/pPy) tract in the proximal 36-bp region (–85 to −50), which is essential for VEGF promoter activity^19^(**Figure S1**). This domain provides 3 potential Sp1 binding sites and a part of thedomain can participate in the formation of higher order G4 structure **(Figure 1)**. Surprisingly this domain shares similarity with our earlier researched cMYC gene promoter.

**Figure 1:**
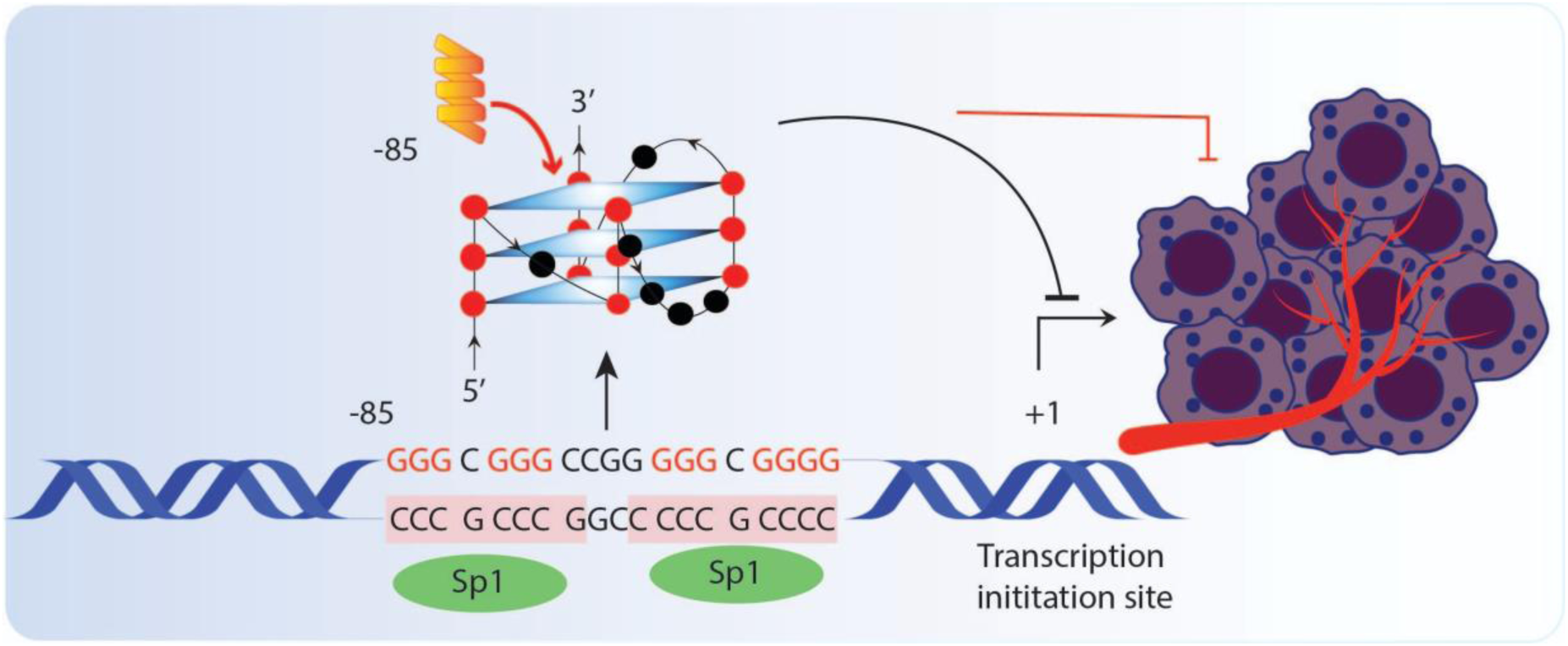
The Polypurine/polypyrimidine (pPu/pPy) tract in the proximal 36-bp region (–85 to −50) in VEGFA promoter which can shuffle between transcriptional inhibitor G-quadruplex and transcriptional activator duplex structure. Peptide targeting and stabilizing the VEGF G-quadruplex thus lowering transcription and inhibiting angiogenesis.

The G4 domain overlaps with two potential Sp1 binding site. This zone has also been revealed to possess highly dynamic conformation and shuffle between G4 and duplex structure. The G4 structure, which is further stabilized by K^+^ ions, can inhibit the VEGF-A transcription. During cancer progression the VEGF-A G4 succumbs to cellular pressure and fails to maintain the stable structure. This shifts the balance to form duplex structure thereby increasing the rate of transcription. Research related to VEGF-A G4 suggests that the transcription of the gene can be manipulated by ligand-mediated G-quadruplex stabilization^20,21^. But off target effects, toxicity and lack of selectivity limit the use of small molecules/ligands as a clinical tool. Nevertheless, anti-cancer bioactive peptides (ACPs) demonstrated a huge potential in G4 targeted therapeutic applications, making them a good prospect for theranostic agents. In therapeutic applications, ACPs have already been found advantageous due to higher on-target specificity, selectivity, biocompatibility, lower accumulation in tissues, lower immunogenicity, lower off-target effects, and reduced toxicity than conventional anticancer therapeutics^22,23^. Due to these criteria, specially designed G4 selective anti-cancer peptides can shine light on the shortcomings of other non-selective G4 targeted therapeutics. It’s imperative to consider that preclinical and clinical research on VEGF-A targeting has demonstrated promise. This reinforces the feasibility of targeting Vascular Endothelial Growth Factor (VEGF) exclusively, resulting in a comprehensive strategy envisioned to impede the pathway of cell survival and commence an apoptotic cascade. The amalgamation of previous research emphatically advocates for the proposition that VEGF-A G4 represents a highly promising target for anti-cancer therapy.

In the present study we wanted to derive a peptide that can stabilize VEGF-A quadruplex with high specificity and prevent unfolding to inhibit VEGF-A expression in cancer cells. We incorporated a similar hypothesis as mentioned in our previous studies related to peptide designing. Additionally, we used bioinformatic algorithms to understand the biochemical and biophysical characteristics. Based on the parameters, the peptide library consisting of RK15, GK15, RK9 and KW10, peptides were successfully synthesized by solid state peptide synthesis. The cytotoxicity and the apoptosis of the peptides towards cancer cells was observed with respect to MCF10A human breast epithelial cell line. To check if peptides could provide additional stability to the quadruplex, Circular Dichroism spectroscopy and Isothermal titration calorimetrywas used. Further to correlate the thermodynamic and kinetic properties of binding profiles of the peptide with its structural aspects, an atomic level structural investigation was performed by structure calculated from solution Nuclear Magnetic Resonance (NMR) and Molecular dynamic (MD) simulations. Angiogenesis is a dynamic biological process characterized by the degradation of the vascular basement membrane, proliferation, and migration of endothelial cells, subsequent formation of a lumen structure, and the establishment of a mature vascular network^24^. The intricacies of angiogenesis involve the precise regulation of diverse cell types, including endothelial cells, smooth muscle cells, and inflammatory cells. Notably, vascular endothelial cells demonstrate a remarkable capability for rapid proliferation, migration, and differentiation in response to physiological stimuli^25^. VEGF has the ability to activate numerous downstream pathways, such as the PI3K-Akt/mTORC2 route, the PLC-MEK pathway, and the Src-FAK pathway that are responsible for vascular permeability, cellular proliferation and cellular motility^26–28^. Through qPCR we have checked the expression of these VEGF pathway genes upon stabilization of VEGF quadruplex by de novo designed peptides.

Additionally, the overall expression profile of VEGF-A and MAPK by Western blot, under treated and non-treated condition, gave us definitive picture of our designed peptide’s ability to control VEGF expression. Once we determined the best candidate for transcription control, we tried to reduce the peptide length to make a bare minimum small peptide for quadruplex binding. For this we developed RK9 and KW10 peptides to check the effect on VEGF transcription.

## 2. Result and Discussion

### 2.1. Design of peptide

Through analyzing the placement of the loop residues and distance of Guanines of VEGF quadruplex we understood that π-π interaction will play a mighty role in stabilizing the topology. For this amino acid with benzene group would be an ideal candidate. With this logic we enforced placement of Phe and Trp at regular interval for RK15 **(Table 1).** We also used the latest random forest (RF)-, support vector machine (SVM)-, and eXtreme gradient boosting (XGBoost)-based algorithms to predict the active Cell-penetrating anticancer peptides (Cp-ACPs). RK15 was predicted to be Cp-ACP by all three algorithms. As a control peptide we designed GK15 by mutating 1 amino acid to ensure rigor and robustness of the algorithm. GK15 was deemed non-Cp-ACP by all the 3 algorithms. In the next phase while reducing the peptide length we followed the same protocol and designed KW10 and RK9 by analysing the RK15-G4 interaction point (detailed later in the chapter). RK9 was interpreted as Cp-ACP by all three algorithms while KW10 was regarded as non-Cp-ACP by SVM and XGBoost but Cp-ACP by RF-based algorithm. KW10 acted as a control peptide to RK9. In every case the model was built based on two independent cell-penetrating peptide (CPP) and anticancer peptide (ACP) subclassifiers by blending numerous compositional and physiochemical-based elements using the multi-layered recursive feature elimination (RFE) method. The result also pointed that the algorithms applied worked perfectly as both the control peptide GK15 and KW10 acted poorly in lending anticancer effect on the cell while RK15 and RK9 had high success in inducing apoptosis in cancer cells.

**Table 1:**
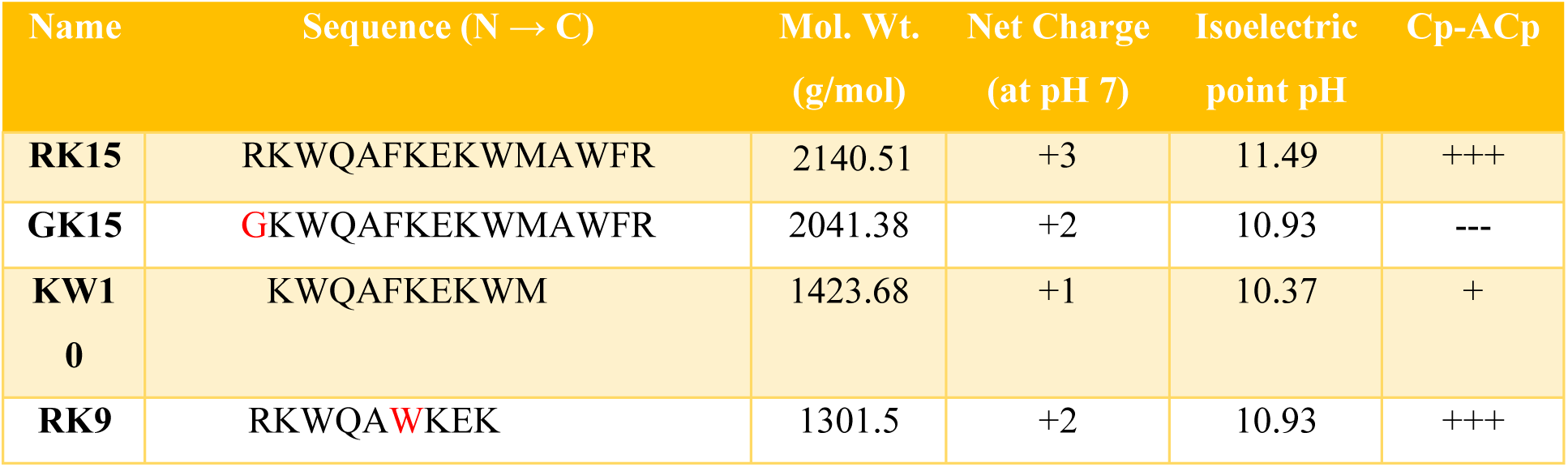
Physiochemical properties of synthesized peptides.

### 2.2. Determination of thermodynamics and structural stability VEGF-A quadruplex upon peptide binding

The peptides RK15 and GK15 were subjected to Isothermal titration calorimetry for decoding their efficacy towards VEGF G4. GK15 revealed no interaction at all with the said G4 while RK15 divulged very strong interaction (K_d_ = 3.1 µM). The strong interaction might be attributed to the placement of Arg at the terminus which has already been proven in the earlier chapter to induce DNA phosphate backbone binding. The thermodynamics revealed that the interaction is enthalpy driven **(Figure 2A).** It can be seen that VEGF G4-RK15 interaction result from enthalpy-driven adsorption processes with a loss of entropy (ΔH < 0 & ΔS < 0) relating to a general prevalence of van der Waals interactions, electrostatics and hydrogen bond formation^29^ **(Figure 2A).** As DNA is negatively charged the Electrostatic interactions will make positively charged RK15 anchor to its backbone. A relatively high –TΔS indicates that RK15 affinity is also supported by hydrophobic interaction. The insertion of peptide into the reaction cell in an aqueous medium breaks the hydrogen bonds between water molecules.

**Figure 2:**
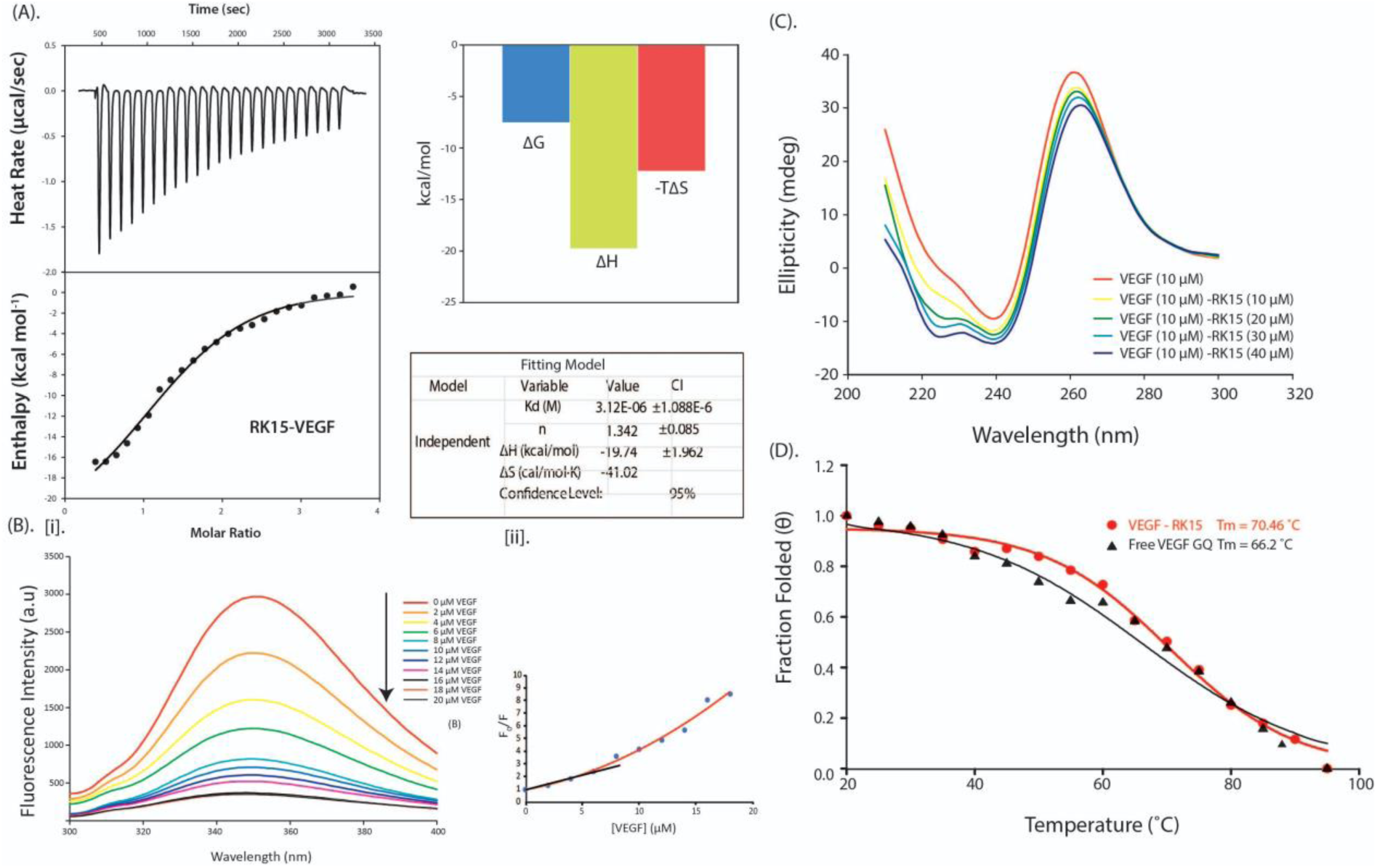
[A]. Isothermal titration calorimetry (ITC) profile of RK15 with VEGFA promoter quadruplex. Left Figure denotes the thermogram. The top panel displays the isothermal plot of RK15-VEGF G4 complex formation, whereas lower panel represents the integrated binding isotherm generated from the integration of peak area as a function of molar ratio. The solid line represents the best fit data using ‘independent binding model’. The binding follows an Enthalpy driven reaction (top right profile), which tends to be propelled by hydrogen-bonding and van der Waals interactions. The top right figure denotes the contribution of Enthalpy and Entropy in RK15-VEGF G4 interaction. Bottom Right denotes the thermodynamic parameters. [B].K15 binding to human VEGFA G4**. (i)** Dose-responsive quenching of RK15 (5 μM) fluorescence emission spectrum with increasing concentration of VEGF G4. **(ii)** Plots of F_0_/F as a function of [VEGF]. F and F_0_ represent the RK15-quadruplex fluorescence intensity at 351 nm with and without quadruplex DNA, respectively. All the samples were measured in 10 mM potassium phosphate buffer (pH 7.0) containing 100 mM KCl. λex = 280 nm[C]. Far UV CD spectra of VEGFA G4 with increasing concentration of RK15.**[D].** CD melting profile of free VEGFA G4 and VEGFA G4 in complex with RK15. All experiments were carried out using 10 mM potassium phosphate buffer containing 100 mM potassium chloride at pH 7.0.

Water molecules that are contorted by the presence of the RK15-VEGF complex generally participate in a network of new hydrogen bonds and form ice-like cage structure called a clathrate cage. This orientation makes the system more structured with a decrease of the total entropy of the system. Hydrophobic interactions are relatively stronger than other weak intermolecular forces. To further authenticate the ITC result we carried fluorescence titration. Peptide-G4 interaction is accompanied with either augmentation or quelling of the fluorescence intensity of the peptide. Change in fluorescence intensity in a concentration-dependent manner can be used to obtain the binding constant and stoichiometry for the RK15–VEGF G4 interaction. Excitation of RK15 at 280 nm yielded an emission spectrum characterized by an intense peak at 350/351 nm **[Figure 2B.(i)].** Fluorometric titration with increasing concentration of pre-annealed VEGF G4 stemmed in the quenching of RK15 fluorescence for the peak at 350/351 nm suggesting a robust interaction **[Figure 2B.(i)]**. Fluorescence intensity was plotted against DNA concentration and was fitted by using the standard Stern–Volmer (SV) equation. Surprisingly, the SV plot from this quenching analysis revealed a linear relationship at lower concentration and a nonlinear upward regression on higher concentration. Such a non-linear SV plot signifies a heterogenous fluorescing system which is intensified by formation of static complexes with DNA. Considering the lower concentration plot, the calculated quenching constant (K_SV_) was found to be 2.2X10^6^ M^-1^.

However, the SV plot with a deviation from linearity is also a common observation for the mixed static/dynamic quenching and efficient singlet exciton migration. But the importance of this strong interaction can only be fulfilled if the peptide can successfully stabilize the VEGF G4. For this we employed Circular dichroism spectroscopy. CD spectroscopy can be used to decipher not only the stability of the VEGF G4 but also the structural changes, if any, upon peptide binding. Titration of RK15 upon VEGF G4 resulted in slight change in the ellipticity of the VEGF G4 which depicts minor structural perturbation **[Figure 2(C)]**. This slender change and fluorescence data suggests that the peptide rests on top of the VEGF quartet and binds to it disrupting the natural chemical structure. We further heated the complex from 20 C - 95 C to analyse the thermal denaturation. As seen in the [**Figure 2(D)]**, free VEGF G4 had a T_m_ = 66.2 C but the melting temperature of RK15-VEGF was increased by ∼ 4 C (T_m_ = 70.46 C). It is well known that quadruplexes with single nucleotide loop are thermodynamically more stable than sequences having two or more nucleotides in their loops^30^. VEGF G4 falls into the 2^nd^ category. Here increase in T_M_ evidently implies the complex is thermodynamically more stable than that of the free VEGF G4. Melting of VEGF in presence of GK15 revealed a comparable melting temperature (ΔT_m_ = 1.8 C), which further points that GK15 has failed to stabilize the VEGF G4. This further confirms the ITC data which indicates no interaction between the said peptide and quadruplex.

### 2.3. Investigation of atomic level interaction of peptide-G4 complex

The interaction of the peptide RK15 with DNA was explored by performing NMR titrations. On titration of VEGF Quadruplex gradually with increasing concentrations of the compound, it was observed that Guanines 4, 5, 9, 15, 16 and 18 showed distinct chemical shifts and changes in peak structures **(Figure 3A).** This indicates that the RK15 binding to quadruplex structure involves the G4, G5, G9, G15, G16 and G18 bases. Unfortunately, upon successive titration all G4 imino signals experience significant line broadening due to dynamic exchange processes upon continued πpeptide titration. As a result, the newly formed complex resonances could not be analyzed and assigned to a particular topology. Next to get detailed information of complex formation at the atomic level molecular dynamics simulation was performed. For docking studies, we modelled the RK15 in PEP-FOLD3 server. It offers a faster coarse-grained approach for de novo linear peptide native like conformation prediction in solution.^31,32^ The VEGF G4 (PDB ID: 2M27) structure was downloaded from RCSB PDB site. The docking was carried out in HDOCK.^33^ HDOCK uses hybrid docking algorithm and free docking protocol for protein-protein and protein-DNA/RNA docking.^34,35^ The model was selected based on very low docking score of −223.57 kcal/mol which in turn indicated the strong interaction between molecules. The complex was solvated with TIP3P waterbox in amber to mimic the cellular chemical environment. Since this "default" geometry may not correspond to the actual minima in the ff14SB force field and may also result in conflicts and overlaps with atoms in other residues, it is always a good idea to minimize the structure. The solvated complex was further minimized with ff14SB forcefield in two steps. Next the system was relaxed and equilibrated for 10 picosecond and simulated for 1 ns. MD simulation showed insights of mechanism of binding of RK15-VEGF G4 interaction.^36,37^ The result shows that RK15 locks onto VEGF G4 using strong polar contact between Lys2-G16 and Lys7-G9 at one terminal **(Figure 3C).** On the other terminal, indole group of Trp13 makes several non-covalent perpendicular stacking interactions with G5, G21 and T22 **(Figure 3B)**. Among all the aromatic rings containing group tryptophan offers strongest binding ability because of best π-electron donating capability. Strategically placed Trp10 also facilitates van der waals and electrostatic connections between G5 and G9. Interface analysis reveals that Trp3 follows the same suit and interacts with G16 and G9. The other connections are listed in Table 2. The aromatic groups, specially the tryptophan with no direct hydrogen bond formation between both rings suggests the partial π-electron transfer of the indole ring to the unoccupied orbitals of guanine base even in its ground state. Thus, this π-π charge-transfer force could be thought of as being a major factor stabilizing the binding between the indole ring and guanine base. The aromatic group containing residues (viz. TRP and PHE) took the leading role in stabilizing RK15-VEGF G4 complex. The simulation results successfully mimicked the NMR spectroscopy data where we see a change in the G5, G16 and G9 peak characteristics.We have also studied NMR titrations with the mutated VEGF-G4 sequence in which G12,G13 residues are mutated T12,T13(**Figure S4**).

**Figure 3.**
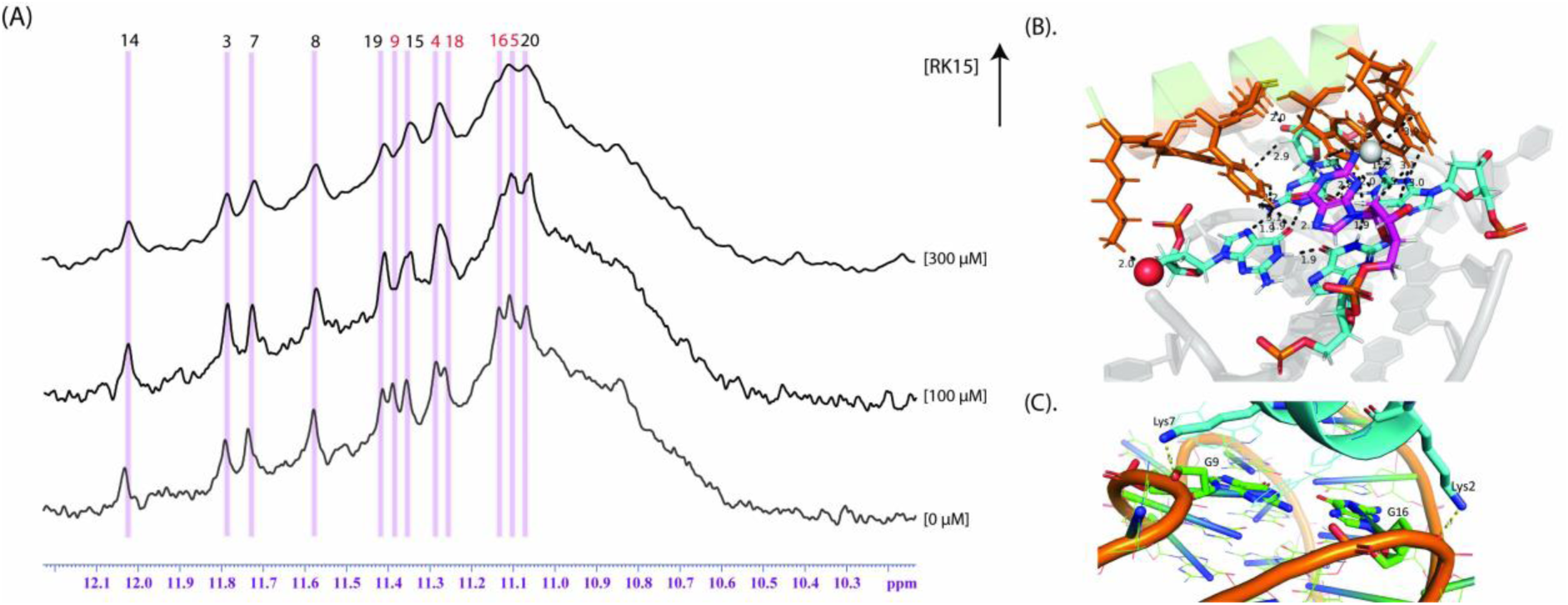
**(A).** 1D proton NMR spectra of VEGFA (300µM) titrated with increasing concentration of RK15 (0-300 µM). The number of imino resonances in the downfield region 10.0–12.0 ppm corresponded to the number of G-tetrad-associated guanines of VEGFA quadruplex. 1D ^1^H NMR was recorded using Bruker Pulse program ‘zgesgp’ with a spectral width of 24 ppm, number of scans of 256, and calibrated pulse length of 12.93 μsec. The red number signifies the Guanines that are participating in Interactions. **(B)** Important polar and non-polar Interactions between RK15 and VEGF G4.(C).Through strong polar contact RK15 locks into VEGF G4 between Lys2-G16 and Lys7-G9 at one terminal.

### 2.4. Shortening of the RK15 peptide increases the potency of interaction

The peptides are more effective if the size is smaller as they can evade degradation by proteases and peptidases. Moreover, any foreign body has the tendency to incite immune response but lower the molecular weight lower is the chance of eliciting an immune response. Since RK15 is a 15 amino acid long peptide we next thought to decrease the size by selecting the key residue like W for interaction and develop the core area as the minimum binding domain. We developed two peptides-RK9 and KW10- and checked their binding potential to VEGF G4. We implemented ITC to check their binding pattern and thermodynamics of interaction. While RK9 disclosed similar strong interaction (Kd = 5.1 µM) as RK15 **(Figure 4A)**, KW10 initiated a no binding at all with VEGF G4. However, the thermodynamics signalled a much higher contribution by hydrogen bond formation (ΔH_RK9_ < ΔH_RK15_) **(Figure 4A. ii)**. Analogous -TΔS (albeit a little higher) like RK15 indicates similar chemical interaction between RK9 and VEGF. Fluorescence emission spectra was next performed to unveil the quenching with increasing concentration of VEGF G4. Excitation of RK9 at 280 nm yielded an emission spectrum characterized by an intense peak at 350/351 nm **(Figure 4B).** Like the RK15 pattern, fluorometric titration with increasing concentration of pre-annealed DNA stemmed in the quenching of RK9 fluorescence for the peak at 350/351 nm suggestinga robust interaction.

**Figure 4.**
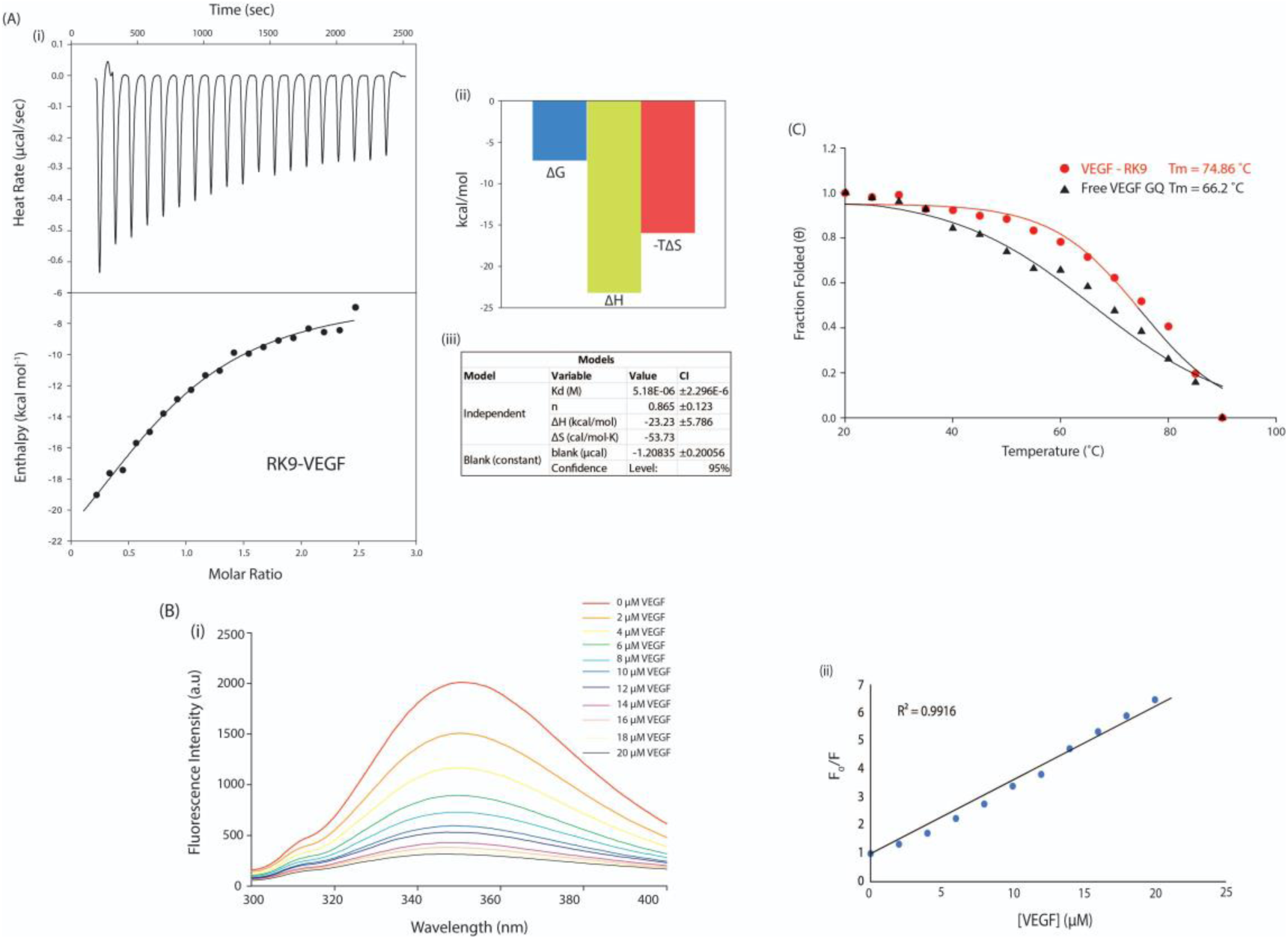
Biophysical characterization of RK9. **(A).** Isothermal titration calorimetry (ITC) profile of RK9 with VEGFA promoter quadruplex. (i)The top panel displays the isothermal plot of RK9-VEGF G4 complex formation, whereas lower panel represent the integrated binding isotherm generated from the integration of peak area as a function of molar ratio. The solid line represents the best fit data using ‘independent binding model’. (ii) The binding follows an Enthalpy driven reaction, which tends to be propelled by hydrogen-bonding and van der Waals interactions. (iii) The thermodynamic profile of the Interaction. **(B).** Fluorescence spectroscopic study of RK9 binding to human VEGFA G4. (A) Dose-responsive quenching of RK9 (5 μM) fluorescence emission spectrum with increasing concentration of VEGF G4. B) Plots of F0/F as a function of [VEGF]. F and F0 represent the RK9-quadruplex fluorescence intensity at 351 nm with and without quadruplex DNA, respectively. All the samples were measured in a 10 mM potassium phosphate buffer (pH 7.0) containing 100 mM KCl. λ_ex_ = 280 nm**. (C).** CD melting profile of free VEGFA G4 and VEGFA G4 in complex with RK9. The presence of RK9 increased the Tm by 9 C. All experiments were carried out using 10 mM potassium phosphate buffer containing 100 mM potassium chloride at pH 7.0.

Fluorescence intensity was plotted against DNA concentration and was fitted by using the standard Stern–Volmer (SV) equation. Surprisingly, the SV plot from this quenching analysis revealed a linear relationship. The linearity of plot suggests that only one type of quenching ensues in the system^38^. The K_SV_ value acquired from the slope of the linear plot was found to be 0.2X10^6^ M^-1^. Such a large value implies presence of strong contact among RK9 and VEGF G4. This strong binding ideally should stabilize the VEGF G4. We employed thermal melting by CD spectroscopy to check the stability offered by RK9 to VEGF G4. As seen in Figure **4C**, free VEGF G4 had a melting temperature of 66.2 C but in presence of RK9 the T_M_ of VEGF G4 was increased by ∼9 C (T_M_ = 74.86 C). Compared to RK15, RK9 almost doubled the melting temperature. Data from ITC, Fluorescence spectroscopy and CD Spectroscopy evidently demonstrates that shortening of RK15 leads to higher quadruplex stability without affecting the binding affinity.

### 2.4. Anticancer efficacy and downregulation of oncogenic expression by the peptides

We reasoned that since RK15 and RK9 both are binding and stabilizing the promoter G4 with high efficiency, the expression level of VEGFA should be negatively affected and will propel the cancer cells towards apoptosis^39,40^. In this study, the peptides had been taken up for the evaluation of the cellular fate of the MDA-MB-231 cells. To test the cytotoxicity of the peptide and cell viability upon its treatment we performed XTT assay. RK15 showed the highest anti-neoplastic activity with IC_50_ value of 8.97 µM. GK15 and KW10 as predicted, initiated no/low cytotoxicity. RK9 yielded an IC_50_ value of 11.58 µM which is likeRK15 (**Figure 5A**). Further flow cytometric studies using Annexin-V-FITC/PI stainingsuggested that RK15 and RK9 sensitized almost 71.26% and 70.56% MDA-MB-231 breast cancer cells to apoptotic death whereas GK15 could push mere 6.6% of the cell towards apoptosis **(Figure 5B)**. The results showed that there was no significant change with GK15 and KW10 treatment can be observed. **(Figure S6)**To shed more light on the mode of action, we next checked if the peptides are altering the VEGFA expression. On the other hand, earlier research suggests that p42/44 MAPKs protect the cells from apoptosis and promote angiogenesis as it is principally linked with cell proliferation and differentiation.^41–44^ Based on the results of the western blot analysis, it can be inferred that GK15 and KW10 treatment has no discernible effect on the expression of either VEGF or MAPK. There was no significant change of expression with the increasing concentration of GK15. However, on treatment with RK15, there was a noticeable decrease in the expression of both VEGFA and MAPK. Following increasing RK15 treatment of 20 µM and 40 µM, there is ∼ 45% and ∼50% reduction in p42/44 MAPK expression (**Figure 5C**). This supports our earlier apoptosis data which shows RK15 to promote apoptosis.

**Figure 5:**
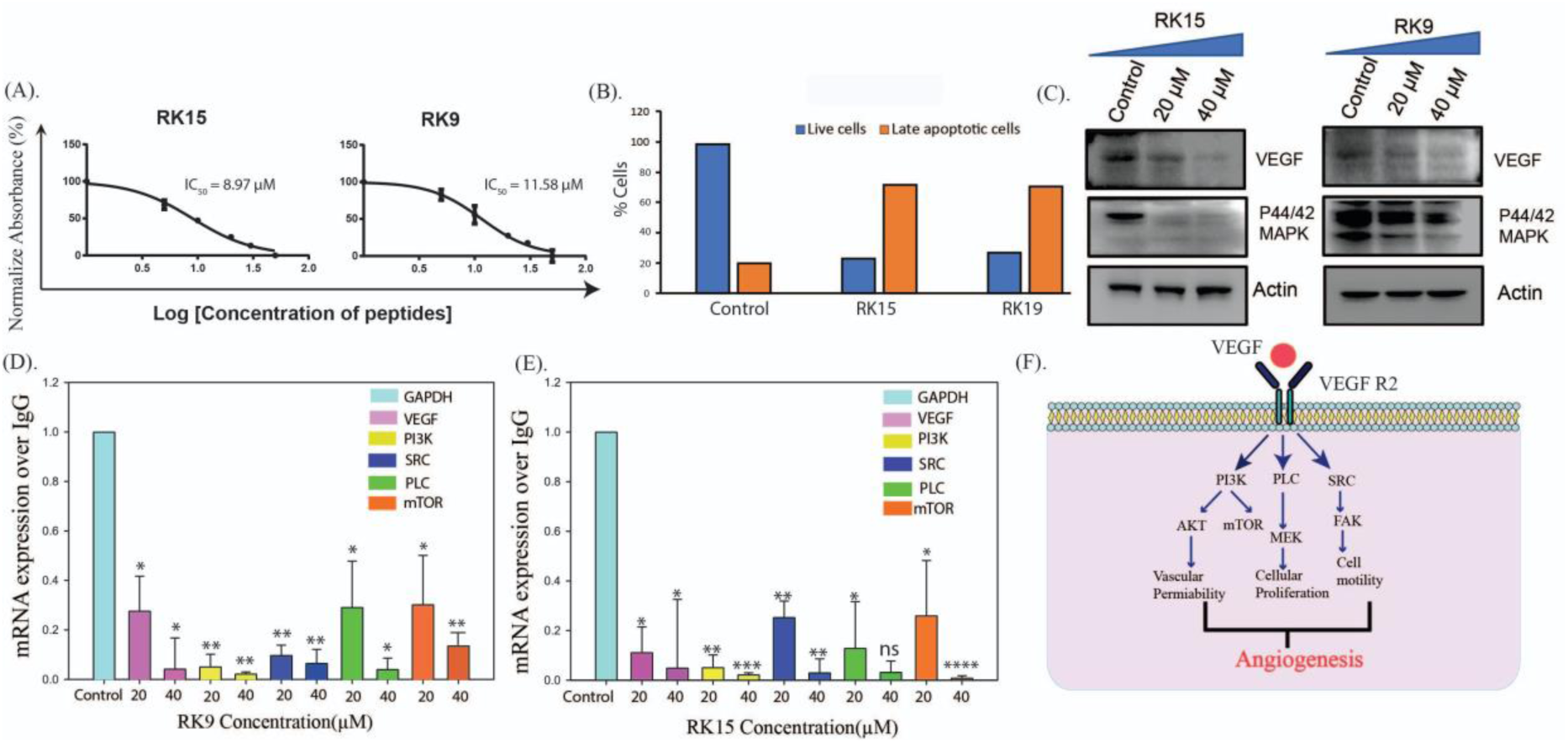
(A). Cytotoxicity of the RK15 and RK9 measured by XTT assay. (B). Percentage of cell viability of MDA-MB-231 cells treated with peptides, RK15 and RK9 after 24 hours incubation. (C) Western blot analyses of the VEGFA and p44/42 MAPK expression in MDAMB-231 cells treated with RK15 and RK9. (D). We treated the MDAMB-231 cell line with two different concentrations of RK9, 20µM and 40µM, in order to investigate the expression of genes associated with the VEGF pathway that are involved in angiogenesis. The qPCR analysis showed that the expressions of VEGF, PI3K, SRC, PLC, and mTOR decreased significantly as the concentration of RK9 increased. (E). We treated the MDAMB-231 cell line with 10µM and 25µM RK15 to assess the expression of genes related to angiogenesis associated with the VEGF pathway. The gene expression was analyzed using GAPDH as a reference. As the concentration increased, the levels of VEGF, PI3K, SRC, PLC, and mTOR showed a significant decrease.

This suggests that RK15 treatment may operate as an angiogenesis pathway inhibitor and apoptosis promoter, leading to a decrease in P42/44 MAPK expression as the activation of p42/44MAPKs significantly enhances VEGF gene expression. RK9 was also found to follow the footprint of RK15. Upon its treatment with 40 µM there was ∼74% and 35% decrease in the expression of VEGF and P42/44 MAPK respectively. We have treated the MDAMB-231 cell line with 20µM and 40µM RK9 and RK15 to observe the expression of VEGF pathway-related genes responsible for angiogenesis. Subsequent analysis of gene expression was performed against GAPDH. The expressions of VEGF, PI3K, SRC, PLC, and mTOR decreased significantly with increasing concentration in both treatments of RK9, RK15 (**Figure 5D**). The decrease in expression was more pronounced in the case of RK9.

The CAM model is a highly effective and widely used approach for assessing the angiogenic potential of purified components and intact cells. The CAM is created by the union of the mesodermal layers of two developmental structures: the Allantois and the Chorion of the avian embryo. CAM, as a highly vascularized tissue and being immune compromised, is an excellent indication of the anti-angiogenic or pro-angiogenic activities of test substances. After identifying RK15 as the most efficient peptide, we conducted a CAM experiment to evaluate its effectiveness as an anti-cancer treatment agent. The 72-hour-old, fertilized egg, at t=0, was carefully broken to preserve the integrity of the embryo. The experimental requirement was met by treating with a concentration of 60 µM RK 15. No treatment was administered during the control setup. A manual countdown of the vessels formed after treatment reveals that the treated egg produces fewer vessels than the control **(Figure S8).**

### 2.5. Shortening of the peptide reduced the specificity to VEGF G4

G quadruplex of various oncogenes share similar structural features and hence it is difficult to develop oncogene specific ligand. If a ligand binds to multiple G4 then it becomes difficult to inhibit specific oncogenic expression. Hence, to conclude if RK15 and RK9 were VEGFA specific we employed ITC to measure their interactions with various promoter G4 structures. We titrated the peptides with ZEB1, cMYC, and KRAS G4 structures to determinespecificity. ITC result revealed that RK15 bears no binding at all with any quadruplex apart from VEGFA but RK9 divulged binding to multiple structures **(Figure 6A).** Interaction with KRAS, ZEB1 and cMYC revealed multiple sites (2 sites), sequential two site binding and independent two site binding respectively **(Table S1).** Important observation here is that RK9 demonstrated 2:1 bonding with all the above-mentioned parallel G4. This is because, RK9 is small sized peptide and structurally these parallel G4 has enough space to accommodate two of such peptides. ΔG value suggests interaction of RK9 with KRAS was strongest followed by cMYC and ZEB1. This proves that though RK9 shows higher stabilization to VEGF G4 it lacks specificity. To understand the atomic nitty gritty of why RK9 is showing lack of specificity with VEGF, we conducted docking and molecular dynamic simulation. Simulation revealed that RK9 is docking to the 3’ end of the VEGF G4 and Lys2 is mediatingthree strong H-bonds with G16 and G20. The sleek non structured RK9 fitted itself into the top of the quartet and formed non-covalent interaction with G15-16, C17 and G18-20. The close placement (within 3.5 Å) of Trp3 allows it to form strong electrostatic interaction with G16, C17 and G19 **(Figure 6B).** RK15 was able to bind to C6 (present in the loop) and G5 with Phe14 and Trp13 respectively present on the other loop. Comparing with RK15 it is evident that the large surface area of RK15 allowed it to bind to multiple residues thereby increasing its specificity, whereas RK9 was able to bind to only selected few residues very strongly which increased the potency of contact but compromised its specificity. This data also corroborates our finding that RK9 binds to other said G4 in 2:1 ratio.

**Figure 6.**
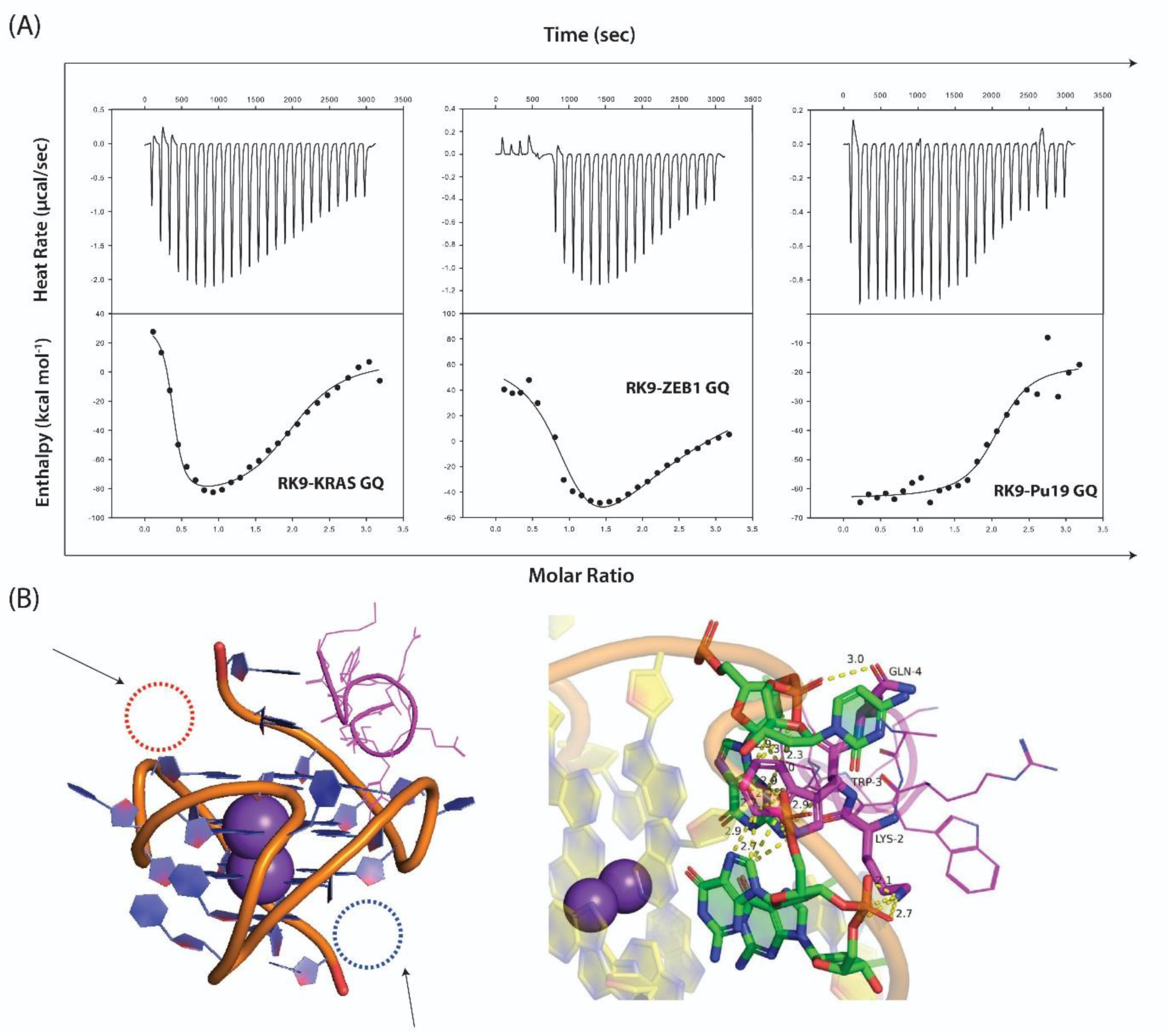
**(A).** Isothermal titration calorimetry (ITC) profile of RK9 with various promoter quadruplex. The top panel displays the isothermal plot of RK9-VEGF G4 complex formation, whereas lower panel represents the integrated binding isotherm generated from the integration of peak area as a function of molar ratio. The solid line represents the best fit data using ‘independent binding model’. RK9 shows interaction with all these parallel G quadruplexes. **(B)** (left) Model of RK9-VEGFA complex. The red and blue circle are other potential binding domains. The long structure of RK15 made it possible to extend to zone marked with red to interact with the residues present there. (right) Important polar and non-polar Interactions between RK9 and VEGF G4.

There is a staunch difference in the way RK15 and RK9 interacts to VEGF G4. RK15 was seen to sit on the top of the 3′- quartet while interacting with the quartet G5/9/16. RK15 uses the loop residues to clasp the entire G4 surface by tethering to the three loops (G21, T22 in one loop, C10 in the diagonally opposite loop and C6 in the adjacent loops) **(Figure 7A)**. RK9 on the other hand forces itself into the 3′ loop and contacts guanines of all the three quartets. The placement of Trp3 in RK9 and RK15 contrasts a lot **(Figure 7B)**. RK9 tandemly interacts with every residue at the extreme 3′- tail from G15-T22 leaving the other part of G4 free **(Figure 7D)**. Trp3 in RK9 plays a major role in forming non-polar contacts with a guanine of every quartet. Apart from Arg41, Ala5 and Glu8 every other residue is participating in polar or non-polar interaction with VEGF. Therefore, we could see an elevated ΔTm in case of RK9 as it is holding and stabilizing 3 quartet plane which is absent in RK15. But again, since RK9 is binding majorly to guanines of quartets it is lacking specificity to a single quadruplex. RK15 might be binding to a single quartet but it encompasses a larger surface area while tethering to two diagonal loops. This explains its low ΔTm but high specificity. The strategically positioned amino acids (K3W4, F6K7, K9W10 and W13F14) are leading the main interactions with G4 **(Figure 7C)**. This demonstrates that peptide must be of a standard length to establish unique binding nature.

**Figure 7.**
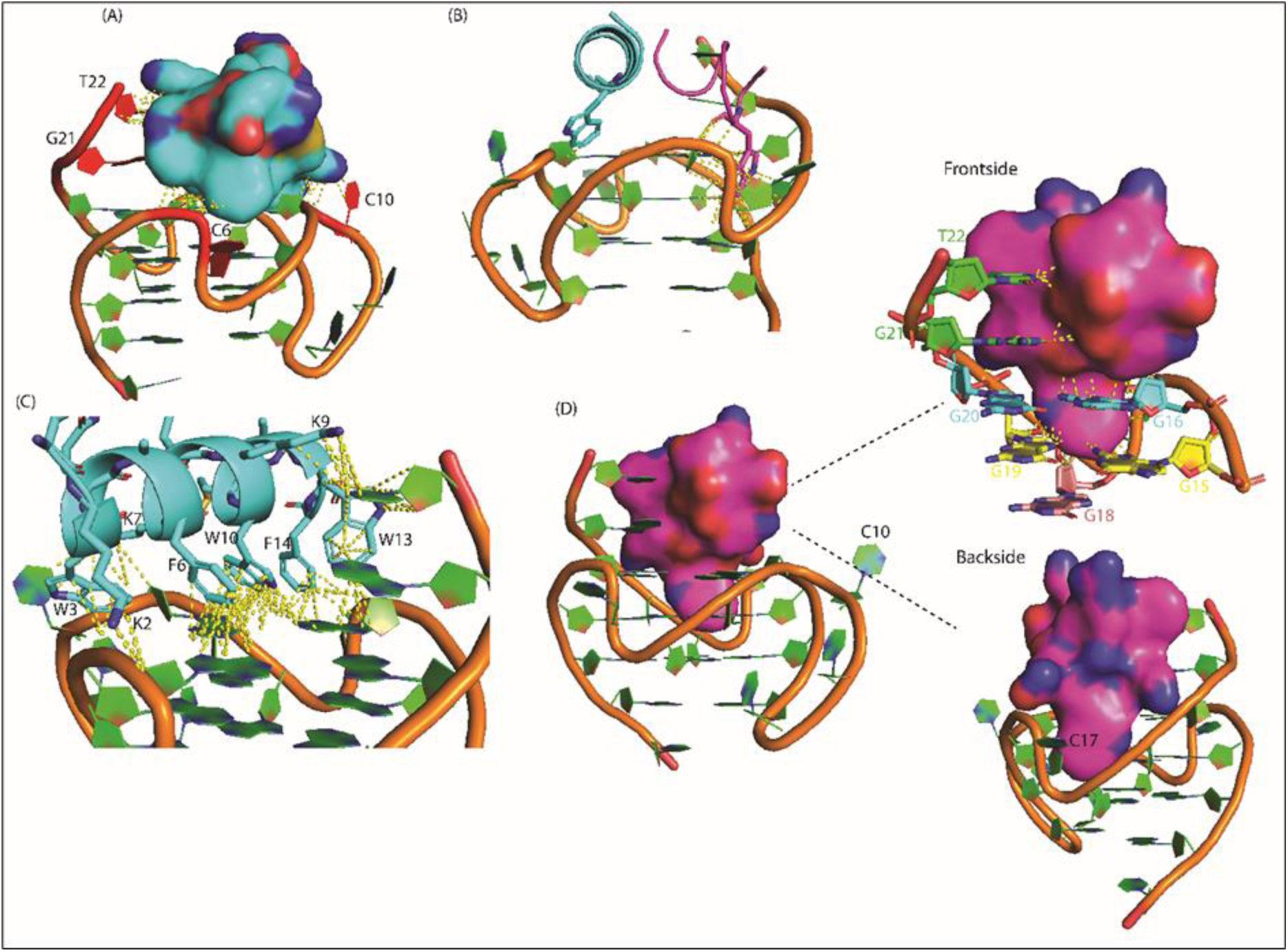
Differences in the binding of RK15 and RK9 with VEGF G4. (A). RK15 binding to the 3′ quartet of VEGF G4. Red nucleotides indicate loop residues in diagonally opposite strand. (B). Difference in the position of Trp3 in RK15 (blue) and RK9(magenta). Trp3 in RK9 is protruding through the loop to interact with guanines of all the quartets while in RK15 it is interacting with C10 and G16. (C). All polar and nonpolar interactions of RK15 majorly formed by Trp, Phe and Lys residues. (D). Binding of RK9 with the 3′- tail of VEGF G4. RK9 interacts with guanines of all 3 quartets. Wheat indicates 5′ quartet. Yellow signifies middle quartet. Blue indicates 3′- quartet. Green signifies 3′ loop residues.

## 3. Conclusion

In this study we have exhibited the development of peptide ligands specific to VEGFA G4. The peptides were mainly constructed by utilizing the SVM, RF, and XGBoost, algorithm which independently distinguish the anticancer efficacy and plasma membrane transducing active sequences from their nonactive equivalents. We developed an active molecule and its non-active analogue as a control. The peptides developed by this mechanism acted in par with the prediction. Peptides offer a unique perspective as they contain dynamicity and conformational flexibility of a protein but cell membrane penetrability of a small molecule. The low toxicity, and greater surface area allows the peptide to selectively chelate to specific G4s. A significant enhancement to the peptide helicity content makes sure that the peptides can easily move past the cell membrane. Angiogenesis encompasses multiple downstream signaling pathways that regulate cellular motility, vascular permeability, and proliferation. The downregulation of VEGF pathway gene expression observed after treatment with RK9 and RK15 suggests that peptide therapy may represent the next generation of therapeutic agents. Though both RK9 and RK15 reduced the expression of VEGF-A signalling pathway, RK9 showed a more pronounced effect in reducing angiogenesis. *De-novo* designed peptide RK15 demonstrated strong binding to the VEGF G4 while offering significant stability. Curtailing the structure of RK15 elevated the ΔT_m_ of VEGFA G4 significantly but compromised its G4 specificity. This indicated that having a substantial surface area allows the peptide to make several interactions throughout the quadruplex landscape thereby making it more specific. Peptides act as a unique ligand because being neither small molecule nor protein, they happen to be at the nexus of these two classes of therapeutics. Due to our limited resource, we were unable to generate libraries of peptides for further studies, but we believe that peptidomimetics using these sequence-based studies can lead to the development of peptides which are excellent complement or even preferable alternative to small molecule and biological therapeutics.

## 4. Materials and Methods

### 4.1. Peptide design and Synthesis

The anti-cancer property and cell penetrating ability of the peptides, using the random forest (RF)-, support vector machine (SVM)-, and eXtreme gradient boosting (XGBoost)-based algorithms, were checked by CpACpP server (http://cbb1.ut.ac.ir/CpACpP/Index). The physicochemical properties were analyzed in silico through pepcalc server. The designed peptides were synthesized using a Solid-phase **p**eptide synthesizer (AAPPTEC Endeavor 90) following Fmoc strategy. Fmoc-protected amino acids were procured from Novabiochem and sequentially coupled from the peptide’s carboxyl-terminal, followed by Fmoc deprotection using 20% piperidine solution for 60 and 30min, respectively. N, N-diisopropylethylamine (DIPEA) was used as an activator base, and benzotriazol-1-yl-oxytripyrrolidinophosphonium hexafluorophosphate (PyBOP) was used as an activator in dry-Dimethylformamide (DMF) as the solvent. The peptide was synthesized in 0.05 mmol synthesis scale using Rink Amide MBHA resin. After washing in DMF, the peptide-attached resin was cleaved by standard resin cleavage cocktail solution containing 92.5% TFA, 2.5% Milli-Q water, 2.5% TIS, and 2.5% phenol. These cleaved filtrates are then precipitated using diethyl ether as the solvent, dissolved in water and acetonitrile and purified with reverse-phase HPLC system (SHIMADZU, Japan) with Phenomenix C18 column using Acetonitrile and HPLC-grade water as dual solvent system **(Figure S3)**. The purified peptide was then concentrated in rotavaporizer and lyophilized overnight. To validate the correct peptide synthesis, peptide masses were measured by MALDI-TOF mass spectrometry with α-hydroxycinnamonic acid matrix (Bruker Daltonics flex Analysis) and correlated with its expected mass (Figure S5).

### 4.2. Circular Dichroism

The experiments were carried out in Jasco-J815 CD Spectrophotometer with Peltier cell holder, and temperature controller CDF-426 L at 25 °C. Thermal unfolding experiments were done for peptide-bound complexes to study whether the peptides stabilize VEGF G-quadruplex. First, the peptides were titrated into 10μM of putative quadruplexes, then samples were heated from 20 to 95°C having the temperature gradient of 5, delay time of 150 seconds, scan speed of 100 nm/min, step size of 1nm and a bandwidth of 1nm. The measurements were carried out using a standard quartz cuvette having path length of 1mm and reaction volume of 350 µl. Since the quadruplexes unfold at higher temperatures and revert to their initial conformations upon renaturation, a ‘two-state transition’ model (folded and unfolded) was assumed to analyze the melting curves and estimate the melting temperatures (Tm) by fitting the data points into sigmoidal curves. Under heat-induced denaturation, molar ellipticity value is minimal at 20 d e g C and maximal at 95 deg C. Each spectrum was accumulated after 5 min of incubation to ensure stable complex formation between G-quadruplexes and peptides. The plot of fraction folded vs temperature was fit with thesigmoidal three parameter equation:

### 4.3. Fluorescence spectroscopy

Fluorescence titration was performed using Hitachi spectrophotometer (F-7000 FL spectrophotometer). Fluorescence spectra were recorded using a 1 cm cuvette. The excitation wavelength was fixed at 280 nm and the emission wavelength was scanned from 300 nm to 400 nm. To determine the mode of binding of peptides with quadruplex sequences, 10 μM of peptides were taken and titrated with increasing concentration of VEGF G4. The Stern-volmer equation was plotted and the K_SV_ were determined as described earlier^45^.

### 4.4. Isothermal Titration Calorimetry

The thermodynamic Attributes of interaction between peptide and putative DNA G4 was studied using Affinity ITC, TA instruments at 25 deg C. The designed synthetic peptides and oligonucleotide sequences were prepared in 10mM potassium phosphate buffer with 100mM KCl. Samples were extensively vacuum degassed for 10 min prior to performing the experiment. The DNA G-quadruplex sequences (405µM) taken in the syringe was titrated against the peptides taken in the cell (10µM), to eject 2µl every 120sec. A control setup consisting of oligonucleotides interacting with the buffer (without any peptide) was used. Equilibrium association constant (Ka), enthalpy (ΔH) and entropy (ΔS) of association were obtained after fitting each isotherm using the NanoAnalyze v3.10.0 software from TA instruments. Δ*G* was calculated from the thermodynamic parameters using the following equation:

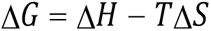

### 4.5. 1D and 2D ^1^H–^1^H NMR spectroscopy and Structure Elucidation

The synthesized peptide RK15, GK15, KW10 and RK9 were confirmed by NMR spectroscopy in Bruker AVANCE III 700MHz and 500MHz NMR spectrometer equipped with 5mm SMART probe. The experiment was carried out in an NMR tube with an active sample volume of 600μl. Peptide of concentration 1mM was added to H2O + D20 solvent in 10:1 ratio with TSP (3-(trimethylsilyl)-2,2,3,3-tetradeuteropropionic acid) which was used as the internal standard. 1D ^1^H spectrum was taken with ‘zgesgp’ Bruker pulse program with 64 scans, spectral width of 20.52 ppm, acquisition time of 1.14 sec and calibrated pulse length of 7.2 μsec. The presence of sharp NMR peaks at the region 8.5-7.5 confirmed the presence of peptides in the sample. Consequently, the sample was titrated with LPS until line broadening was observed. LPS was added to mimic the cell membrane so that the peptide can take up the native conformation **(Figure S2)**. 2D ^1^H-^1^H Total correlation spectroscopy (TOCSY) (to analyze through bond interaction) and 1H-1H Nuclear Overhauser effect spectroscopy (NOESY) spectra (for analyzing through space interaction) of the LPS-peptide complex was collected with Bruker pulse program ‘mlevesgpph’ with 80 ms mixing time and ‘noesyesgpph’ with 150 ms mixing time at 25°C respectively. NOE intensities were qualitatively categorized as strong, medium, and weak based upon their respective cross-peak intensities from NOESY spectra obtained in presence of LPS. This interproton upper-bound distances were 3.0, 4.0 and 5.0 Å respectively. The lower bound distance was reserved constant at 2.0 Å. To restrict the conformational space for all residues, the backbone dihedral angle (phi) was varied from −30° to −120°. No hydrogen bonding constrains were used for structure calculation. Obtained interactions, sequence, bond length, and angle constraints were used to build the peptide structures using CYANA module. Several rounds of structure refinements were performed andbased upon the NOE violations; the distance restraints were adjusted accordingly. The ten lowest energy structures were selected to generate ensembles of structures of peptides. To understand which Guanine residues of VEGF quadruplex were partaking in interaction with RK15, we slowly titration 300 μM VEGF quadruplex with increasing concentration (0-300μM) of the peptide. Formation of RK15-G-quadruplexes complex was identified by chemical shifts of imino protons in 1H-NMR in H2O/D2O solution. The number of imino resonances in the downfield region 10.0–12.5 ppm corresponded to the number of G-tetrad-associated guanines present in the system. Exchangeable and non-exchangeable protons of Myc22 were observed in the 1D proton spectra under free and CD23-bound conditions using Bruker Pulse program‘zgesgp’with a spectral width of 24 ppm, number of scans of 256, and calibrated pulse length of 12.93 μsec. Finally, the peptide structures were modelled. **(Figure S6)**.

### 4.6. Docking and Molecular Dynamics Simulation

The structure of the peptides and the G-quadruplex (PDB ID: 2M27) were uploaded to online bioinformatics docking program HDOCK, which uses a hybrid algorithm of template-based modelling and ab-initio free docking. The top 10 structures were downloaded, and the structure that has the least global energy was selected. The said docked structure was then solvated with TIP3P waterbox with a dimension of 12 Ǻ in all the three z-, y-, z-dimensions. and neutralized by adding precalculated counter ions. This provides a more realistic environment like solvent environment inside cell systems. The complex was then minimizedin two steps in ff14SB and gaff forcefield. The system was minimized to prevent steric clashes by lowering potential energy to less than 0.001 kcal/ mol/Ǻ. Next the complex was slowly heated to 300 K and equilibrated to relax the system prior to simulation. The structure was finally simulated for 10 ns under NPT conditions.

### 4.7. Western Blotting

MDA-MB-231 cells are seeded in 60 mm dishes and allowed to attach overnight/70% confluency. Then cells are treated with 20 μM, 40 μM peptides. Cells are harvested after 24 hrs of treatment. Whole cell lysates are prepared using the Cell Lysis Buffer (Cell Signalling Technologies) according to the manufacturer’s protocol supplemented with PMSF and Protease Inhibitory Cocktail (Merck). Protein estimation was done using the Bradford’s method using Bradford reagent (Amresco) [1-3]. 40 μg of protein is separated using SDS-PAGE. Then the samples are transferred to polyvinylidene fluoride (PVDF) membranes (Immobilon-P, EMD Millipore) and blocking was done using 5% skimmed milk in Tris-buffered saline (TBS) with Tween 20 (Amresco) for 1 hour at room temperature. Membranes were then incubated overnight at 4°C with the following antibodies: anti-cleaved caspases 3 (rabbit polyclonal, 1:1000 dil. Cell Signalling Technology Inc.), anti-VEGF (mouse monoclonal, 1:500 dil. Santacruz Biotechnology Inc.), cleaved PARP (rabbit polyclonal, 1:2000 dil. Cell Signalling Technology Inc.), anti actin (mouse monoclonal, 1:5000 dil., Sigma Aldrich.) Next day membranes were incubated with peroxidase conjugated anti-rabbit IgG raised in Goat (Sigma, 1:8000 dil.) and peroxidase conjugated anti-mouse IgG raised in Rabbit (Sigma, 1:10,000 dil.) for 2hr at room temperature. Target proteins were visualized on membranes using Clarity TM Western ECL substrate (Bio-rad) and images were captured using Azure 600 system. Image analysis was done using Image J software.

### 4.8. Cell culture

MDAMB231 cell lines were grown in RPMI 1640 media supplemented with 10 % FBS with 70-80% confluency at a density of 1 × 106 cells were taken as 100μl-150μl / well. In another set MCF10A (Normal breast supplemented with 10 % FBS with 70-80% confluency at a density of 1 × 106 cells were taken as 100μl-150μl / well.

### 4.9. Cell viability Assay

Both MCF10A and MDAMB231 cell lines were treated with RK15, RK9, GK15, and KW10 respectively in increasing concentration, i.e. 5μM, 10μM, 20μM, 40μM, 50μM. XTT detection solution was added as per the XTT assay kit protocol (Cell signalling) and placed in the incubator for 2 hours. XTT was converted to a formazan dye due to the activity of mitochondrial enzymes in viable cells. The dye was calorimetrically viewed by spectrophotometric analysis at 450nm, and the lethal concentration at which 50 % of the population was killed was determined by Graph pad prism.

### 4.10. Flow Cytometry

MDAMB231 was cultured in a 6-well plate followed by treatment with the peptides in respective wells at lethal dose concentration determined from XTT assay, for 24 hours. The untreated well was used as control. The flow cytometric assay was performed with BD PharmigenTM Annexin V-FITC Apoptosis detection Kit (Catalog no. 556570) for quantification of apoptosis. The cells were harvested after resuspension in PBS and were subsequently fixed with chilled 80% ethanol added dropwise and stored overnight at −20^0^C. Then the cells were again pelleted down and resuspended in PBS followed by the addition of RNaseA and kept at 37 for 2 hours. Finally, PI was added and kept at 40 for 30 minutes prior to performing the experiment. Data is acquired using a flow cytometer.

### 4.11. Real time qPCR following RNA isolation

The qPCR amplification of genes related to the VEGF pathway and other genes were performed after treatment with RK15 and RK9. The samples for PCR were prepared using the PowerUpTM SYBRTM Green Master Mix (Cat. # A25742), and the primers were designed explicitly for this purpose. The PCR cycles were conducted using the QuantStudio 5 thermal cycler. The thermal cycling conditions were as stated: First step of denaturation: The sample was heated at 95 deg C for 10 minutes. This was followed by 40 cycles of initial denaturation at 95 deg C for 30 seconds, annealing at an acceptable melting temperature for 30 seconds, and extension at 72 deg C for 30 seconds. The ΔΔCt value is calculated by subtracting the target gene’s Ct value from the reference gene’s Ct value (GAPDH). The fold enrichmentis calculated using the formula 2^(−ΔΔCt).

### 4.12. Chorioallantoic membrane (CAM) assay

CAM assay is the most widely used ex-vivo assay used for studying neo-angiogenesis. When examining the angiogenic potential of whole cells and purified components, the CAM model is one of the most effective and widely used methods. To maintain sterility, every experiment was conducted in a cell culture environment with sterile conditions. Before the trials began, all the equipment was sanitized with 70% alcohol. The hatchery provided 2-day-old (48-hour) pathogen-free embryonated eggs, which were stored inside the husks during the journey to the laboratory. The eggs were positioned horizontally in an appropriate egg tray and incubated inside an incubator for one hour at 37°C and 50% humidity without turning. After three days (72 hours), the egg tray with the eggs was removed from the incubator and placed into a laminar flow hood. For the control groups, we used an in-ovo setup in which the eggs were incubated for an additional day without any treatment. For peptide treatment sets, we followed ex-ovo setup in a petri plate to better observe the effect on angiogenesis. Using the relatively sharp edge of a sterile hard cube/rectangle shaped structure that was positioned perpendicular to the horizontal axis of the eggs, the eggs were held in a horizontal position and cracked open. Using a hole puncher, filter papers (thickness 1 mm, diameter 6 mm) are formed into round discs. Before the discs were used as carriers for the peptide solution (RK 15) to the CAM, they were autoclaved and thoroughly dried. Equal amounts of RK15 peptide (60 µM) were absorbed into these circular discs for each of the experimental groups that participated in the angiogenic studies. To effectively observe the effects of thepeptide on angiogenesis, the circular discs were dropped over the CAM precisely at the site of dichotomy of the blood vessels, from where two major blood vessels branched, with theaid of sterilized forceps. After the treatment, the eggs were again placed inside an incubator for the next 24 hours maintaining all the requirements. After 96 hours, egg trays were removed from the incubator and again placed inside laminar hood. A high-resolution digital camera was used to capture images of the embryo and the discs for the CAM assay’s end-point analysis.

