## Supporting files for "In Silico designed cell-penetrating anti-cancer peptide specifically inhibits VEGF-A expression"

**Supplementary Contents:**

1. 1D-$^1$H NMR Spectra of VEGF-Pu22 and VEGF-Pu22 T12 T13
2. 1D-$^1$H NMR titration of RK9
3. 1D-$^1$H NMR titration of VEGF-G4 with RK15
4. Purification of peptide by HPLC and conformation of mass-by-Mass Spectrometry
5. 2D assignment of RK9
6. FACS data (Treatment with KW10, GK15)
7. Chorioallantoic membrane (CAM) assay
8. Realtime PCR Primer
9. Table1: Thermodynamic parameters of RK9 interaction with other promoter G4
10. Table 2: Receptor-ligand interface residue pair(s)

**Experiments performed:**

**1.Preparation of oligonucleotides:**

VEGF-G4 sequence is purchased from Sigma at 1 micromolar synthesis scale in lyophilized form. This sequence is dissolved in 10mM KP and 100mM KCL. These sequences are annealed at 95°C for 5 minutes followed by gradual cooling through overnight which leads to the formation of G-quadruplex in-vitro.

**2.Peptide synthesis:**

The peptides were synthesized using a Solid-phase synthesizer (AAPPTEC Endeavor 90) employing the Fmoc methods. The Fmoc-protected amino acids were obtained from Novabiochem and were linked together in sequence starting with the carboxyl-terminal of the peptide. Then, the Fmoc protecting groups were removed using a 20% piperidine solution for 60 minutes and 30 minutes, respectively. The activator base utilized was N, N-diisopropylethylamine (DIPEA), whereas the activator used was benzotriazol-1-yl-oxytripyrrolidinophosphonium hexafluorophosphate (PyBOP). The solvent used was dry Dimethylformamide (DMF). The peptide was synthesized at a synthesis scale of 0.05 mmol utilizing Rink Amide MBHA resin. Following the washing process with DMF, the resin bound to the peptide was broken down using a standard solution called resin cleavage cocktail. This solution consisted of 92.5% TFA, 2.5% Milli-Q water, 2.5% TIS, and 2.5% phenol. The cleaved filtrates are subsequently precipitated using diethyl ether as the solvent. They are then dissolved in a mixture of water and acetonitrile and purified using a reverse-phase HPLC system (SHIMADZU, Japan) equipped with a Phenomenix C18 column. Acetonitrile and HPLC-grade water are used as the dual solvent system for purification. The isolated peptide was further concentrated using a rota-vaporizer and then subjected to lyophilization for an extended period. To verify the accuracy of the peptide synthesis, the peptide masses were determined using MALDI-TOF mass spectrometry with an α-hydroxycinnamonic acid matrix (Bruker Daltonics flex Analysis) and compared to the anticipated mass[1].

**3.1D-1H NMR spectroscopy and 1D NMR titration:**

Nuclear Magnetic Resonance (NMR) studies were performed on a Bruker 700 MHz spectrometer equipped with a 5mm SMART probe. The samples of VEGF were prepared in a solution containing 10 mM KP-KCL and 10% D2O. For 1D NMR tests, a sample volume of 350 µl was used in a shigemi tube. The spectra obtained from each scan were compared to TSP (3-(trimethylsilyl)-2, 2′, 3, 3′-tetradeuteropropionic acid) as a reference compound with 0.0 ppm. The VEGF-G4 NMR sample was titrated with RK9 and RK15 at a 1:60 (oligo: ligand) ratio. The NMR titration experiments are often performed with VEGF T12 T13.G-quadruplex generated in Pu22-T12T13 as the initial structure and substituted T12 and T13 with the wild-type G12 and G13 residues, which is already reported. We have seen the sharpening of peaks, which indicates the stabilization of the structure followed by destabilization, reflecting the peak's broadening[2].

After the preparation of the RK9 peptide, 1mM of Peptide is subjected to NMR titration. The spectra reveal a broadening of the peak with increasing titration. These results suggest the stabilization of VEGF quadruplex with increasing LPS concentration, and this peptide was then subjected to 2D NMR titration.

**4.Flow cytometry:**

Cell line MDAMB231 was cultured in a 6-well plate and treated with peptides in respective wells at a lethal dose concentration determined from XTT assay. The treatment was given for 24 hours, and an untreated well was used as a control. Quantification of apoptosis was performed using the BD PharmigenTM Annexin V-FITC Apoptosis detection Kit (Catalog no. 556570) through the flow cytometric assay. The cells were harvested and resuspended in PBS and then fixed with chilled 80% ethanol added dropwise and stored overnight at -20°C. The cells were again pelleted down and resuspended in PBS, followed by the addition of RNaseA, and kept at 37°C for 2 hours. Finally, PI was added and kept at 40°C for 30 minutes before performing the experiment. The data was acquired using a flow cytometer[3].

A

### 1.1D-$^1$H NMR Spectra of VEGF-Pu22 and VEGF-Pu22 T12 T13

| | |
|---|---|
| 1.VEGF-Pu22 | CGGGGCGGGCCGGGGGCGGGGT |
| 2.VEGF Pu22 T12 T13 | CGGGGCGGGCCTTGGGCGGGGT |

B.

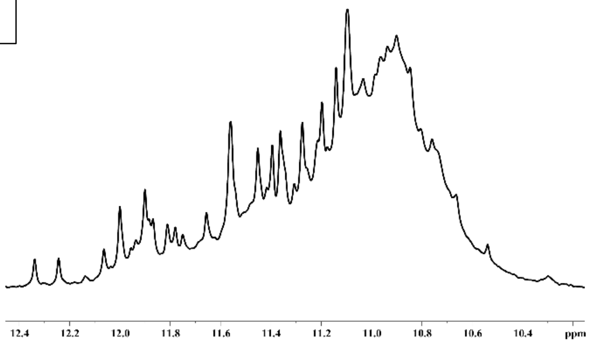

VEGF-G4

C.

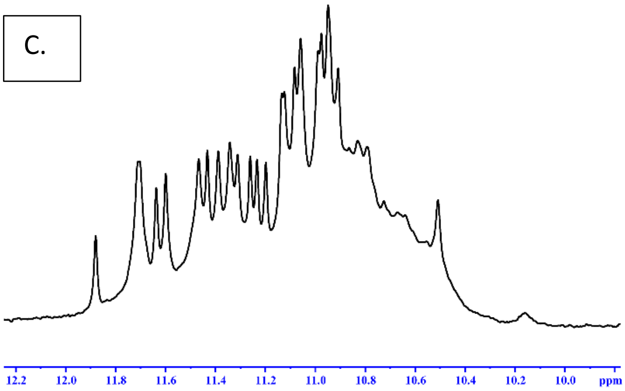

VEGF T12T13

Figure S1: (A).VEGF Pu22 [wild type] and VEGF Pu22 T12 T13[mutated] sequence.(B). The imino region of 1D 1H NMR spectra of the wild-type VEGF-Pu22(C)To create the G-quadruplex in the wild-type VEGF_Pu22 sequence, we used the G-quadruplex generated in Pu22-T12T13 as the initial structure. We substituted the T12 and T13 residues with the wild-type G12 and G13 residues.1D NMR spectra of VEGF-G4 and VEGF T12T13[2]

### 2.1D-$^1$H NMR titration of RK9

a.

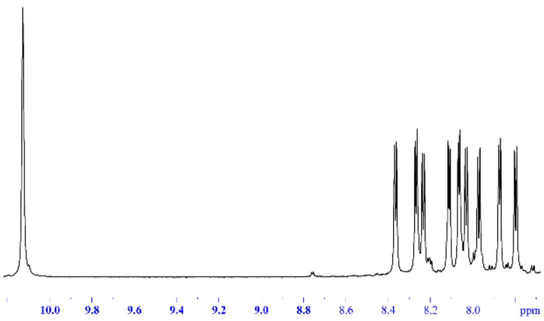

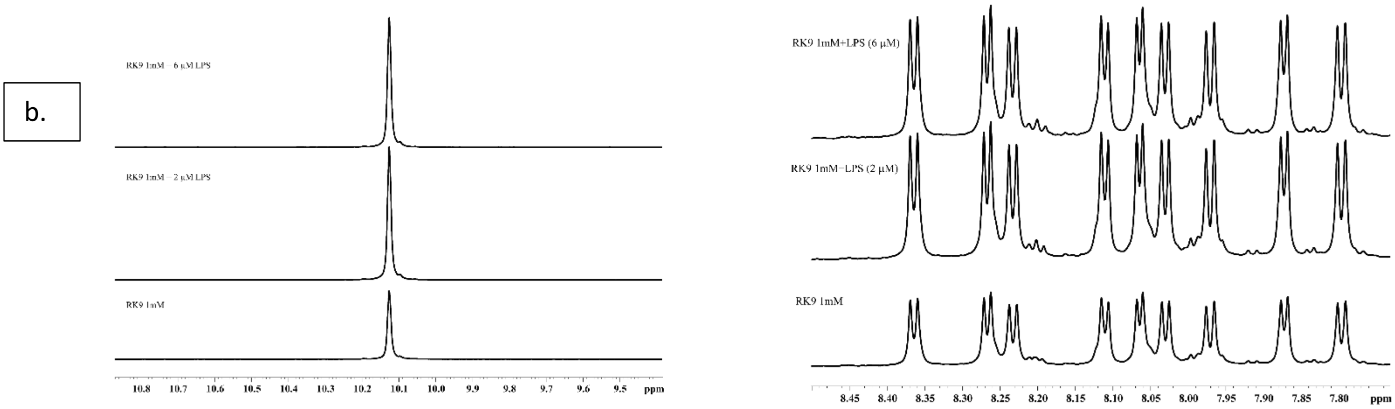

Figure S2: (a). 1D-$^{1}$H NMR spectra of of RK9 peptide.(b).NMR titration of RK9 with increasing concentration of LPS which results sharpening of spectra.

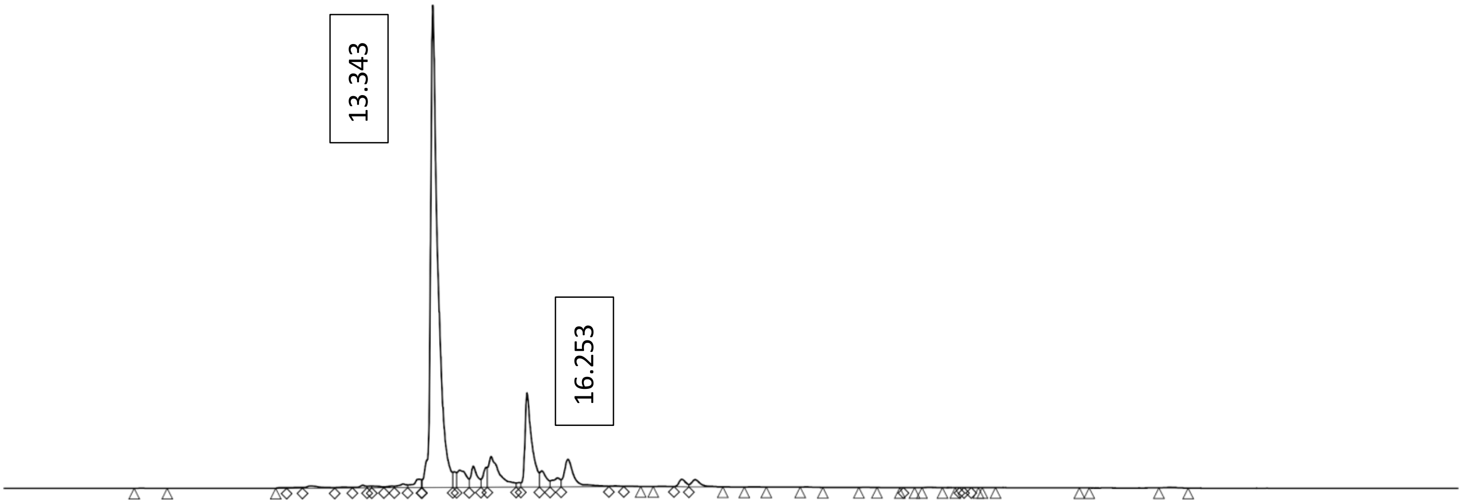

Figure S3: The peak collected during HPLC purification.

| | RT(min) | Area(µV sec) | % Area | Height | % Height |
|---|---|---|---|---|---|
| Peak 1 | 13.343 | 37389724 | 61.39 | 2371671 | 61.05 |
| Peak 2 | 16.253 | 6802896 | 11.17 | 463546 | 11.93 |

**3.1D-1H NMR titration of VEGF-G4 with RK15**

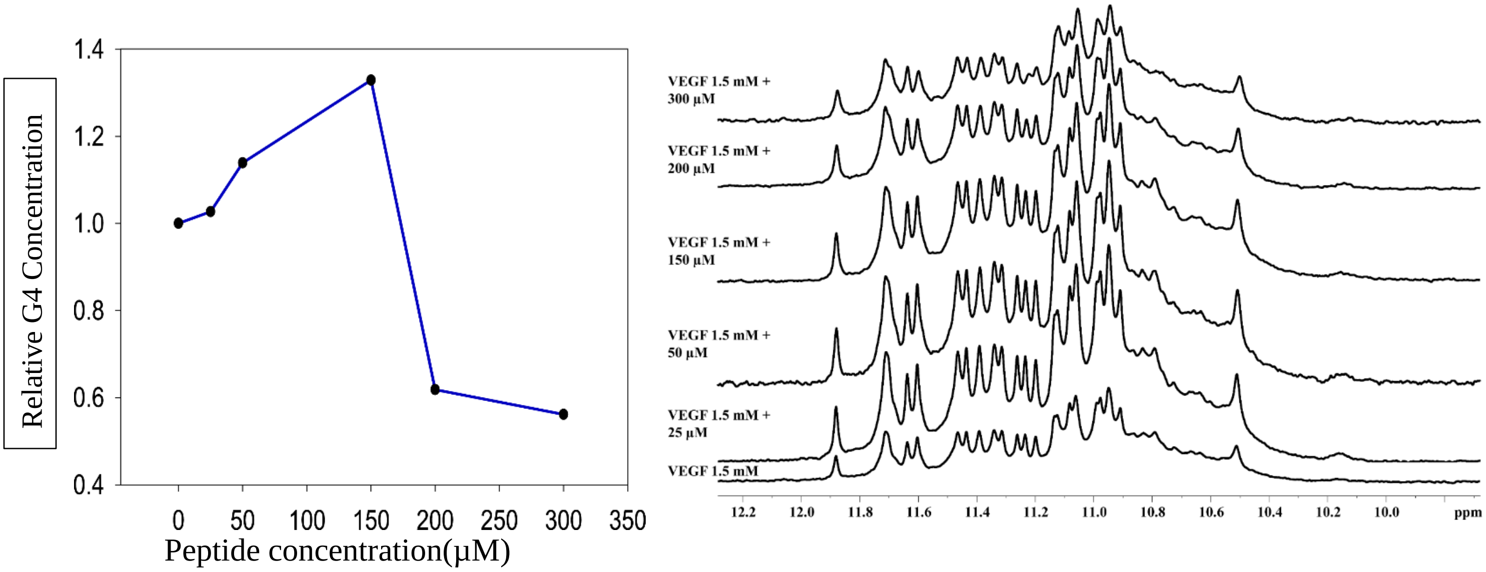

Figure S4: Titration of VEGF T1 T2 with increasing concentration of RK15. With increasing concentration of RK15 the G-quadruplex is stabilized followed by destabilization above 200 μM.

**4.. Purification of peptide by HPLC and conformation of mass by Mass Spectrometry**

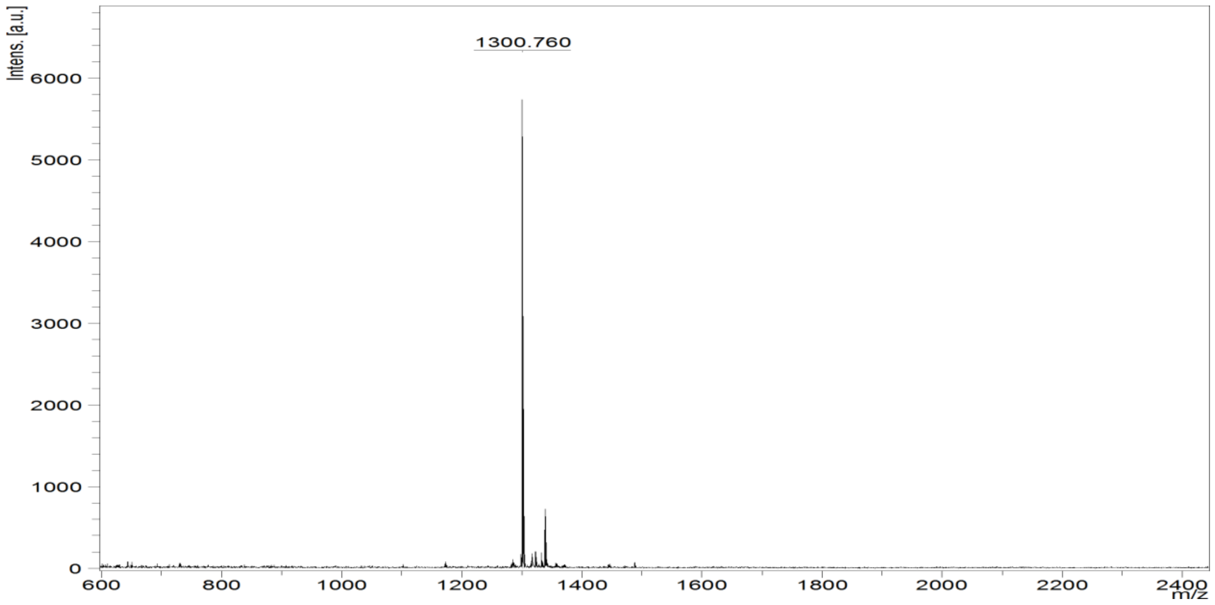

Figure S5. To determine the molecular weight, MALDI-TOF mass spectrometry is performed, which confirms the weight obtained from pepcal.

**5.2D assignment of RK9:**

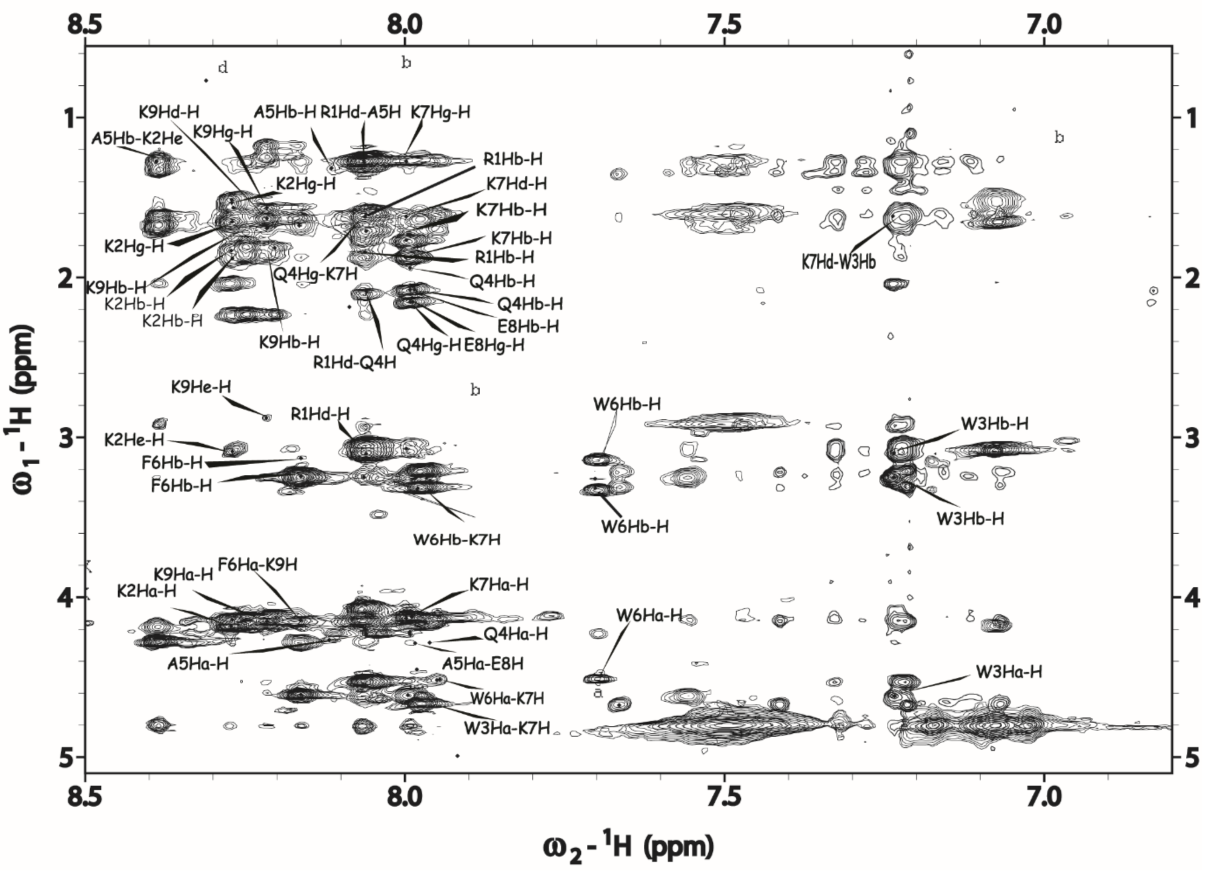

Figure S6:2D assignment of RK9 peptide.

**6.FACS data (Treatment with RK9,RK15,KW10,GK15)**

GK15 Treatment and KW10 Treatment:

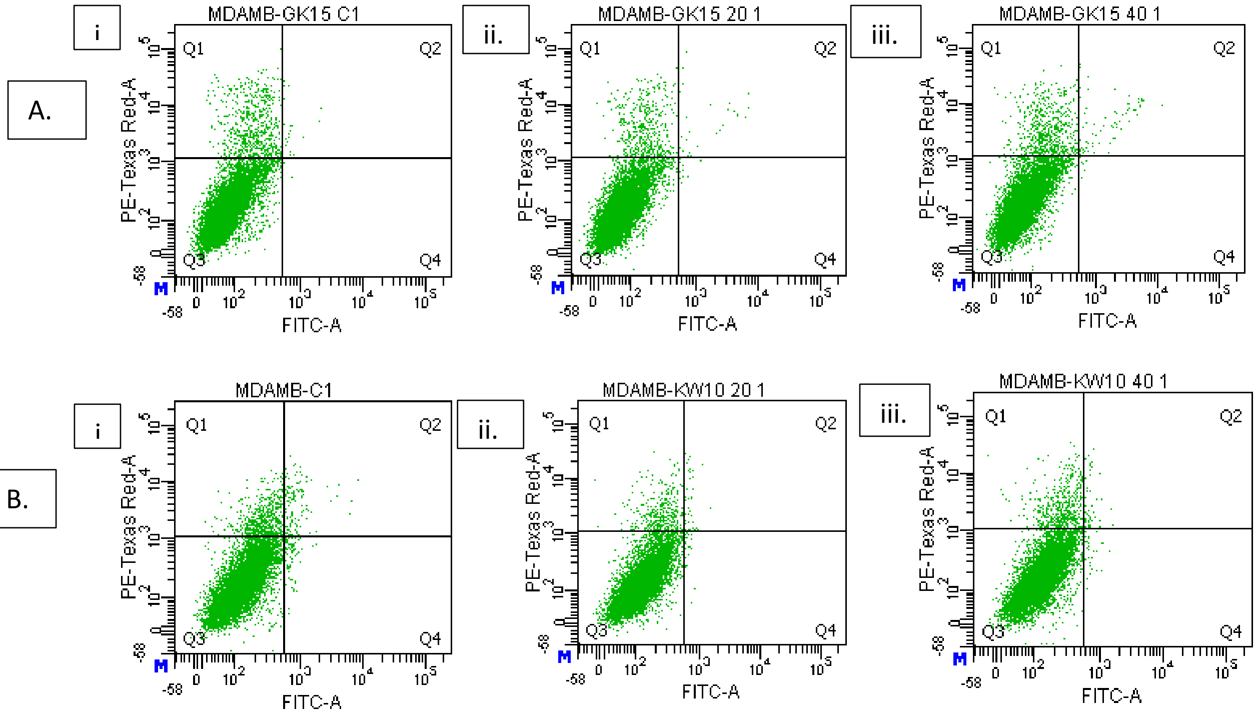

Figure S6: [A]. The control for GK15 treatment is A.i. GK15 C1-. In this experiment, Q1 represents necrotic cells, while Q2 represents late apoptotic cells. In the control experiment, Q1 showed a 6.9% result, indicating no significant change.ii. When treated with 20uM GK15 peptide, MDA-MB-231 cells showed a Q1 5.1%, suggesting no effect with this concentration. iii. However, when treated with 40uM GK15 peptide, the Q1 result was 6.8%, indicating no significant change with this treatment. Q1 5.7%. [B]. KW10 C1- is the control for KW10 treatment in which Q1 6.6% and Q2 1.8% ii.KW10 20 1- MDA-MB-231 cells treated with 20 uM KW10 with Q1 5.5% and Q2 2% .KW10 40 1- MDA-MB-231 cells treated with 20uM KW10 with Q1 5.8%, Q2 0.7%. The results showed that there was no significant change in the treatment as the values of Q1 and Q2 remained almost the same in all the cases. Therefore, it can be concluded that the treatment did not have a significant impact on the cells.

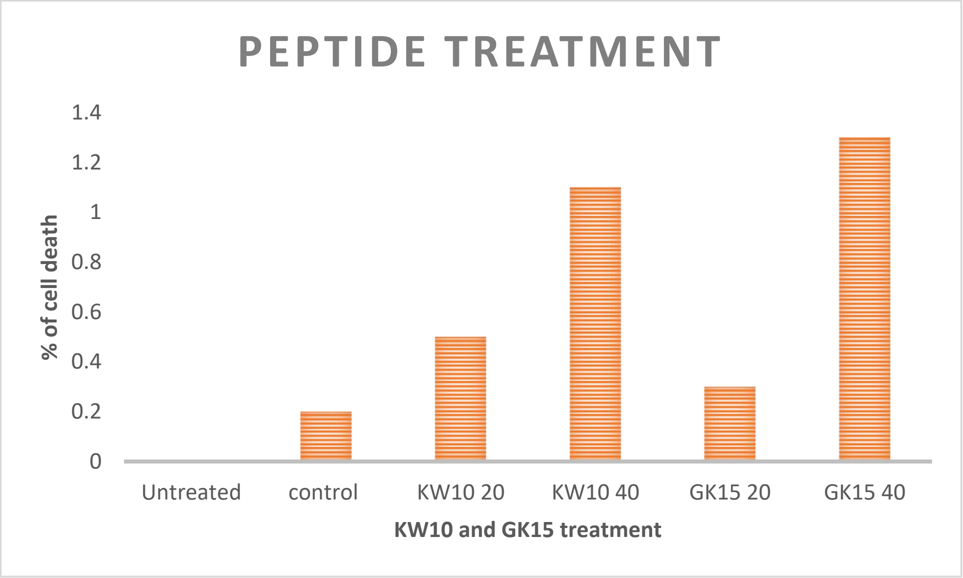

Figure S7: Data representing percentage of cell death against different peptide concentration.

**7.Chorioallantoic membrane (CAM) assay:**

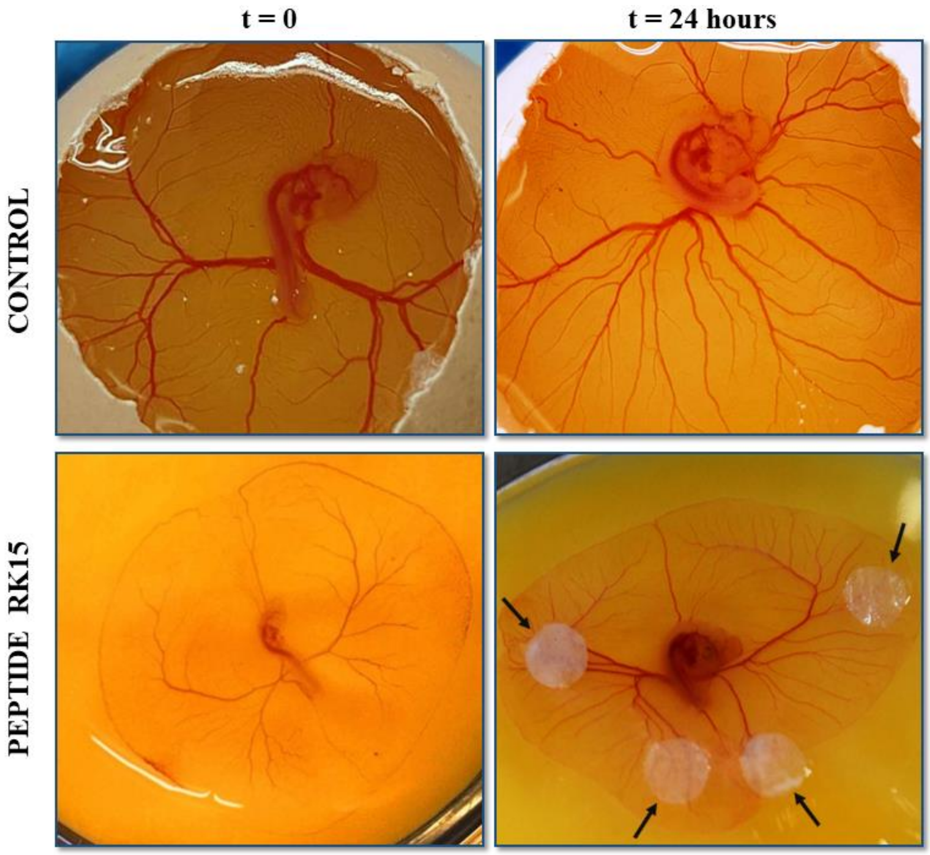

Figure S8: RK 15 inhibits neo-angiogenesis. CAM assay was performed in presence or absence of RK15.The three-day-old (72 hours, t=0) fertilized egg was broken to keep embryo intact. 60 μM RK 15 treatment was done according to experimental requirement. In control set up no treatment was done. After t = 24 hours (96 hours), images were captured to evaluate the formation of new blood vessels in control and RK15 treated sets. Arrow indicates the site of dichotomy of the blood vessels, from where two major blood vessels branched to form new vessels.

### 8.REAL-TIME PCR Primer:

| | |
|---|---|
| SRCFORWARD | AGGTGGAGCACTACCGCATC |
| SRC REVERSE | AGAGTCCGTCGGCATCTGTG |
| PI3K FORWARD | TGCTATGCCTGCTCTGTAGTGGT |
| PI3K REVERSE | GTGTGACATTGAGGGAGTCGTTG |
| PLCγ FORWARD | GGAGCTGAGTATGACAGCACCAAGC |
| PLCγ REVERSE | AGAGGCGGTTGTCTCCATTGACC |
| mTOR FORWARD | ACTGCTTTGAGGTCGCTATGA |
| mTOR REVERSE | TTGCCTTTGGTATTTGTGTCC |
| GAPDH FORWARD | GATGCTGGCGCTGAGTACGTCGTG |
| GAPDH REVERSE | AGTGATGGCATGGACTGTGGTCATGAG |

### 9.Table S1: Thermodynamic parameters of RK9 interaction with other promoter G4 :

| DNA | N (sites) | $K_d$ (M) | ΔH (kJ/mol) | ΔS (J/mol·K) | ΔG (kJ/mol) |
|---|---|---|---|---|---|
| **KRAS** | 0.332 ±0.036 | $1.40 \times 10^{-8}$ | 22.53 ±27.80 | $2.26 \times 10^{2}$ | -44.81 |
| | 1.712 ±0.160 | $1.43 \times 10^{-6}$ | -98.66 ±26.34 | $-2.19 \times 10^{2}$ | -33.39 |
| **ZEB1** | 1 | $9.72 \times 10^{-7}$ $\pm 8.32 \times 10^{-7}$ | -20.76 ±66.72 | 45.5 | -7.2 |
| | 1 | $3.37 \times 10^{-5}$ $\pm 6.95 \times 10^{-5}$ | -547.7 ±999.7 | $-1.75 \times 10^{3}$ | -25.5 |

| | | | | | |
|---|---|---|---|---|---|
| **cMYC** | 2.005 ±0.093 | 4.40 X $10^{-7}$ ±3.34 X $10^{-7}$ | -42.87 ±5.206 | -22.09 | -36.28 |

### 10.Table S2: Receptor-ligand interface residue pair(s):

| VEGF G4 Residue: Chain ID | RK15 Residue: Chain ID | Distance ($A^0$) |
|---|---|---|
| 5A | 10B | 2.143 |
| 5A | 13B | 3.853 |
| 5A | 14B | 3.118 |
| 6A | 14B | 2.827 |
| 8A | 10B | 4.763 |
| 9A | 3B | 4.258 |
| 9A | 6B | 3.038 |
| 9A | 7B | 3.001 |
| 9A | 10B | 2.622 |
| 10A | 3B | 3.027 |
| 10A | 7B | 3.097 |
| 15A | 3B | 4.078 |
| 16A | 2B | 2.998 |
| 16A | 3B | 3.522 |
| 16A | 6B | 2.937 |
| 17A | 2B | 4.267 |
| 21A | 6B | 4.311 |
| 21A | 10B | 4.71 |
